# Large-scale discovery of candidate type VI secretion effectors with antibacterial activity

**DOI:** 10.1101/2021.10.07.463556

**Authors:** Alexander Martin Geller, David Zlotkin, Asaf Levy

## Abstract

Type VI secretion systems (T6SS) are common bacterial contractile injection systems that inject toxic “effector” proteins into neighboring cells. Effector discovery is generally done manually, and computational approaches used for effector discovery depend on genetic linkage to T6SS genes and/or sequence similarity to known effectors. We bioinformatically investigated T6SS in more than 11,832 genomes of Gram negative bacteria. We found that T6SS encoding bacteria are host-associated and pathogenic, enriched in specific human and plant tissues, while depleted in marine, soil, and engineered environments. Analysis of T6SS cores with C-terminal domains (“evolved” cores) showed “evolved” HCP are rare, overwhelmingly encoded in orphan operons, and are largely restricted to *Escherichia*. Using the wealth of data generated from our bioinformatic analysis, we developed two algorithms for large-scale discovery of T6SS effector proteins (T6Es). We experimentally validated ten putative antibacterial T6SS effector proteins and one cognate immunity gene from a diverse species. This study provides a systematic genomic perspective of the role of the T6SS in nature, a thorough analysis of T6E evolution and genomic properties, and discovery of a large number of candidate T6Es using new approaches.

## Introduction

Microbes utilize various antagonistic mechanisms to infect hosts and to kill competing microbes. The type VI secretion system (T6SS) is a membrane-bound, contact-dependent secretion system that injects toxic T6SS effector proteins (T6Es) into neighbouring bacteria and into eukaryotic cells (Cianfanelli, Monlezun, and Coulthurst 2016; Pukatzki et al. 2006). Structurally, the T6SS is a contractile injection system, which shoots a spear-like structure towards neighboring cells. The “spearhead” and “shaft of the spear” are secreted from the attacking cell (Vettiger and Basler 2016). The “spearhead” is made up of a protein called PAAR, and a trimer of VgrG, while the shaft is made up of a column of hollow, donut-like structures, formed from stacks of hexamers of the protein HCP (Jing Wang, Brodmann, and Basler 2019).

T6Es are oftentimes proteins that non-covalently interact with HCP, VgrG, or PAAR, and are thereby secreted into target cells upon T6SS contraction. These are sometimes called “cargo” effectors (Alcoforado Diniz, Liu, and Coulthurst 2015). However, sometimes the HCP, VgrG, and PAAR proteins (“core proteins”) contain an N-terminal core domain, e.g. a VgrG domain, and a C-terminal toxin domain, e.g. an enzymatic actin crosslinking domain. This example of an “evolved” VgrG exists in *Vibrio cholerae* (Pukatzki et al. 2007). Core proteins with C-terminal domains are called “evolved” cores (e.g. “evolved” PAAR). “Evolved” cores are also sometimes referred to as “specialized effectors” (Alcoforado Diniz, Liu, and Coulthurst 2015). Other examples of “evolved” core effectors include VgrG-3 of *Vibrio cholerae*, which has a C-terminal peptidoglycan degrading activity (Brooks et al. 2013), VgrG2B of *Pseudomonas aeruginosa*, which has a C-terminal metallopeptidase activity (Wood et al. 2019), HCP-ET1, which has a C-terminal nuclease domain (Ma, Pan, et al. 2017), and Tse6 of *P. aeruginosa* has an N-terminal PAAR with a C-terminal toxin (Whitney et al. 2015; Quentin et al. 2018).

“Evolved” cores are of interest, yet there has not been a comprehensive study of them. There have, however, been studies of the T6SS operons as a whole bioinformatically. One was published 13 years ago, and used 400 genomes for a first genomic look at the T6SS (Bingle, Bailey, and Pallen 2008). The authors concluded that the T6SS is present in more than a quarter of sequenced bacterial genomes. Furthermore, they showed that the T6SS is mostly present in Proteobacteria (Bingle, Bailey, and Pallen 2008). Another bioinformatic study used 501 prokaryotic genomes to compare and contrast T6SS genes and operons found in these genomes (Boyer et al. 2009). Boyer et al. found that taxonomically, the T6SS is present largely in Proteobacteria. Other bioinformatic studies of the T6SS generally focus on a specific taxonomic subgroup (Repizo et al. 2019; Robinson et al. 2021), and to the best of our knowledge, no broad T6SS bioinformatic analysis has been published recently, despite the precipitous rise of the availability of genomic sequencing data and the growing understanding of T6SS importance in shaping microbial interactions with host and microbe.

T6SS and its T6Es are not only important to microbial ecology (i.e., their highly important role in niche colonization) and pathogen-host interaction (Speare et al. 2018; Logan et al. 2018; Dörr and Blokesch 2018) but also to application as potential new antimicrobials, to medicine and to agriculture. For example, study of T6Es highlights novel bacterial antimicrobial targets. The recent study of T6E Tse8 showed that inhibition of transamidosome activity has antibacterial activity (Nolan et al. 2021). This knowledge can lead to development of new antimicrobials that inhibit the transamidosome, a novel target, to treat infectious diseases. Therefore, it is important scientifically and practically to investigate and discover diverse T6Es. Despite this importance, a recent survey screened Proteobacterial genomes that contain T6SS genes and found that only 42% had effectors of known activity, which suggests a large portion of effectors remain undiscovered or their mechanisms of action are yet unknown (LaCourse et al. 2018).

Generally, T6Es have been discovered in specific model organisms by looking at genetic linkage, i.e. functionally screening genes that are encoded near the T6SS core genes for toxic activity (Lien and Lai 2017). This approach is effective, but is inefficient, and may miss toxins that are not genetically linked to the core components of the T6SS, i.e. “orphan” or “auxilliary” operons, which are far from the T6SS core operon (Crisan et al. 2019). There is also the possibility that there are “super orphan” operons, i.e. T6E genes located in the genomes distantly from any T6SS core genes. Indeed, our group identified one putative T6E family that had no close genetic linkage to known T6SS genes (Levy et al. 2017). Another method to discover T6Es are experimental screens (Lien and Lai 2017), but this method has limited scalability, as it requires growth and mutation of the species of interest.

An alternative method to identify T6Es is to use bioinformatic approaches to mine the wealth of prokaryotic genome sequencing data currently available publicly. Past studies that have used computational approaches to search for new effector genes have relied on algorithms that use 3D protein modelling homology to known enzymatic activities as parameters. The Mougous group has successfully identified a peptidoglycan amidase T6E superfamily using an algorithm that filtered hits for potential amidase activity by protein modelling (Russell et al. 2012), and subsequently used a similar algorithm to search for putative lipase and lysozyme activity, respectively (Whitney et al. 2013; Russell et al. 2013). Another bioinformatic approach used to discover new T6Es was a sequence homology based algorithm, which identified a conserved amino acid sequence motif called the MIX domain that was present in a variety of T6SS effectors (Salomon et al. 2014). A machine learning approach was taken to identify putative T6Es, but the authors did not validate their computational predictions (Jiawei Wang et al. 2018).

A powerful predictive algorithm could be designed that relies on the adaptor proteins of the T6SS (also called “chaperone” proteins). Adaptor proteins in the T6SS context refers to proteins that facilitate the proper folding and loading of their partner toxic effectors onto the T6SS (Unterweger et al. 2015). For example, Tap-1/TEC, an adaptor of a T6SS in *V. cholerae*, was discovered by two separate groups as a chaperone protein that facilitates the loading of a toxic effector onto the T6SS, but is not secreted itself (Unterweger et al. 2015; Unterweger, Kostiuk, and Pukatzki 2017; Liang, Moore, and Wilton 2015). The major families of adaptors studied to date, which contain the conserved domains DUF4123, DUF1795, and DUF2169, all share the feature of being genetically linked to their corresponding effector (Unterweger, Kostiuk, and Pukatzki 2017).

In this study, we perform a broad scale bioinformatic analysis on 11,832 T6SS-encoding genomes, with a special focus on “evolved” HCP, VgrG, and PAAR. We explore the characteristics of “evolved” cores, and we also use this information to discover novel putative T6Es with two distinct algorithmic approaches. We used these algorithms to perform a massive discovery of T6Es in thousands of bacterial genomes and validate their toxicity. We discovered 10 putative T6Es that were found to cause toxicity to *E. coli*, some of which are “super orphans”, i.e. far from any core T6SS genes. Our methodology for large scale prediction and screening is in contrast to the majority of T6SS studies, which mostly focus on discovery of a single effector in a single species, and provides an alternative way for discovery of T6SS effectors.

## Results

### T6SS-encoding genomes make up 19% of all sequenced bacteria

Using conserved domains of the T6SS (Materials and Methods), we found 11,832 genomes with T6SS (Figure 1A, Supplementary Data 1) out of a total 62,075 genomes that we download from the IMG bacterial genome database (I.-M. A. Chen et al. 2019), which represents 19% of the database. Like previous studies (Boyer et al. 2009), we see that the T6SS in our dataset are mainly encoded by Proteobacteria, and overwhelmingly by Gamma- and Betaproteobacteria (85.5% and 11% of all T6SS-encoding genomes respectively; Figure 1B). In contrast, we saw there is a depletion for Alpha-, Delta-, and Epsilonproteobacteria (Figure 1B). Looking at the enrichment of T6SS at the genus level, we see a T6SS enrichment in well-studied Gammaproteobacterial genera, such as *Yersinia*, *Klebsiella*, *Proteus*, *Pseudomonas*, *Vibrio*, *Serratia*, *Escherichia*, *Salmonella* (Figure 1B). Interestingly, the genus *Shigella* seems like an outlier within the Enterobacteriaceae family. Its related genera are T6SS-rich, including *Escherichia*, *Klebsiella*, *Chronobacter*, *Citrobacter*, and *Salmonella*, whereas *Shigella* is for some reason T6SS-poor, and might be using an alternative predominant molecular weapon although T6SS in certain *Shigella* species has been reported (Anderson et al. 2017). The genus with the largest enrichment of T6SS was *Aliivibrio*. This is unsurprising, as it is known that *Aliivibrio fischeri* (*Vibrio fischeri*) uses its T6SS as a critical factor that determines squid light organ niche colonization in a competitive environment (Speare et al. 2018). *Yersinia* also was highly enriched for T6SS, with 96% of sequenced species (428/445) encoding the T6SS marker genes, which is expected given the T6SS’s role in virulence, competition, and ion transport (Yang et al. 2018). We note that our dataset overwhelmingly seems to contain T6SS subtype i, with a handful of Genera known to harbor subtype ii. This is likely because the conserved domains used for search were likely based on subtype i and ii marker domains (Materials and Methods), and likely do not detect more distant subtype iii and iv T6SS.

**Figure 1.**
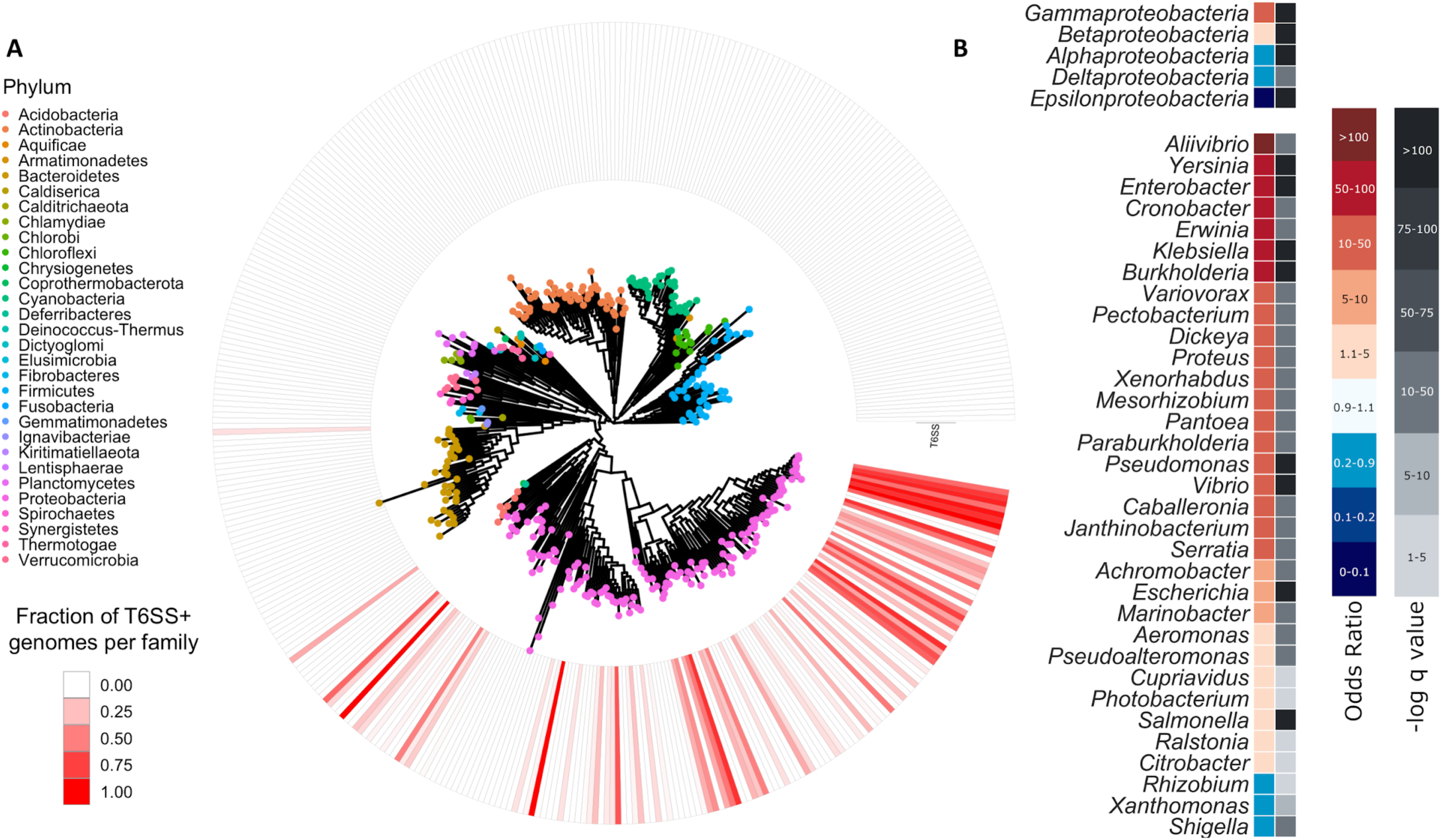
T6SS Genome distribution and enrichment. (A) Phylogenetic tree of taxonomic Families in our database (each leaf represents one Family), with percentage of the members of each Family encoding T6SSs (ring of red shades). Colors on leaves represent Phylum. (B) Statistical enrichment (redder colors) and depletion (bluer colors) of Classes (top) and Genera (bottom) encoding T6SS. Enrichment was calculated using Fisher exact test, and corrected with Benjamini Hochberg method to produce q-value (plotted as -log10 q value in grey).

### T6SS-encoding bacteria live a pathogenic lifestyle and are enriched in specific tissues of humans and plants

We noted that members of these well-studied Genera are of such interest to science in part because they are known for containing pathogenic species. In order to quantitatively understand the lifestyle and environments of T6SS-encoding bacteria, we performed an enrichment analysis of their genome metadata. The metadata includes features about the isolation site and lifestyle of the sequenced microbe. This analysis is statistically corrected for biases in the genomic data (I.-M. A. Chen et al. 2017, 2019; Brynildsrud et al. 2016) (see Materials and Methods). Our analysis showed that T6SS-encoding genomes are indeed mildly enriched for pathogenic lifestyle (odds ratio [OR] = 1.268, q-value = 0.0194; Figure 2; Supplementary Data 2). T6SS-encoding bacteria are enriched in eukaryotic hosts (OR = 4.06); specifically they are enriched in humans (OR = 2.42), cattle (OR = 2.16), and plants (OR = 2.99) (Figure 2). Within humans, T6SS is enriched in microbes isolated from the digestive system (OR = 1.69), the excretory system (OR = 3.16), the urethra (OR = 3.78), and from feces (OR = 5.92). T6SS is, however, depleted from microbes isolated from the reproductive system (OR = 0.09) and from subgingival plaques (OR = 0). In plants, T6SS is strongly enriched in endophytes, which are isolated from the plant interior (OR = 35.5), and from the rhizoplane (root surface) (OR = 5.71). Previous findings showed that T6SS is indeed important for root colonization of certain bacteria (Mosquito et al. 2020). Furthermore, we see T6SS-encoding genomes are statistically depleted from aquatic (OR = 0.218), marine (OR = 0.251), soil (OR = 0.509), and environmental sources generally (OR = 0.264), and also from engineered environments (OR = 0.462), such as activated sludge (OR = 0.164) and bioreactors (OR = 0.09) (Figure 2). Therefore, T6SS can be described as a host-associated microbial toxin secretion system, although not present in all hosts (e.g., T6SS is depleted from bacteria isolated algae). When analyzing the physiological lifestyle of T6SS-encoding bacteria, we observed that T6SS is enriched in motile (OR = 16.7), mesophilic (OR = 3.42), nitrogen fixing (OR = 7.85) bacteria and is depleted from anaerobic bacteria (OR = 0.026) and from bacteria that dwell in radical temperatures. As expected, we detect that T6SS is a feature of Gram negative bacteria and is depleted from Gram positive bacteria (Figure 2, Supplementary Data 2).

**Figure 2.**
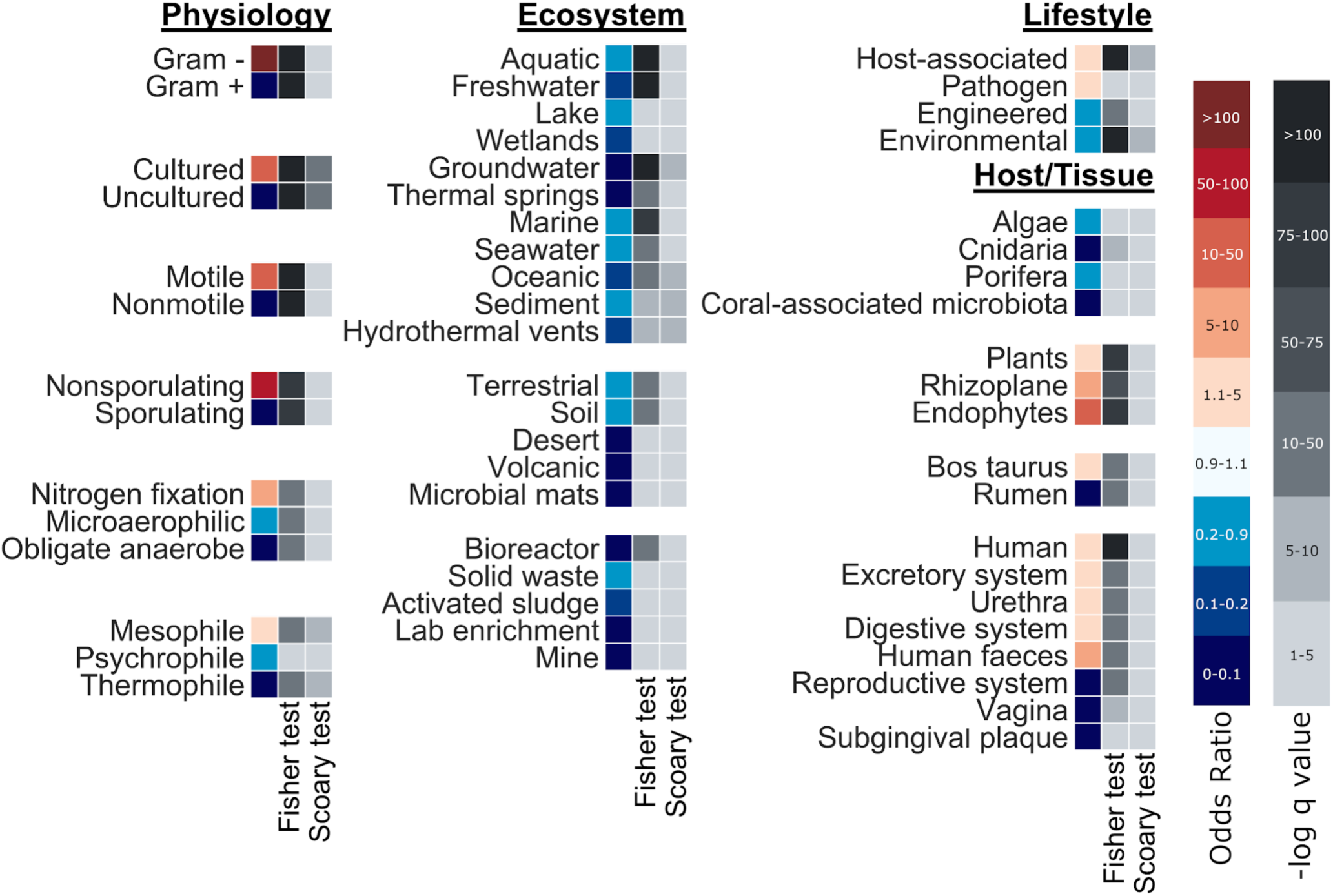
Metadata enrichment of T6SS-encoding genomes. Metadata of T6SS-encoding genomes were analyzed using Scoary (Brynildsrud et al. 2016). The analysis assigns an odds ratio that quantifies the enrichment (redder) or depletion (bluer) of metadata labels using a Fisher exact test. Importantly, Scoary also takes into account phylogeny to correct for taxonomic biases that exist in our genome database. Therefore, we have statistical significance for both the naive Fisher exact test (middle columns) and for the phylogeny-aware Scoary test (right columns), displayed as a -log10 of the q-values.

### “Evolved” HCP are narrowly taxonomically distributed, smaller, and have limited effector repertoires compared to other “evolved” cores

One of the most intriguing aspects of the T6SS are its toxic T6SS effector proteins (T6Es). To be secreted from the attacking cell into neighboring cells, the toxic T6E proteins are non-covalently or covalently attached to the T6SS machinery. The latter are called “evolved” cores. The covalently bound “evolved” T6Es roughly follow a polymorphic toxin architecture, with an N-terminal trafficking domain, and a C-terminal toxin domain (Zhang et al. 2012). In the case of T6SS, the N-terminal trafficking domains are components of the core machinery: VgrG (Spike component), PAAR (Tip of the spike), or HCP (tube component) (Figure 3A). The C-termini many times are enzymes, and we report their overall characteristics in the following paragraphs.

**Figure 3.**
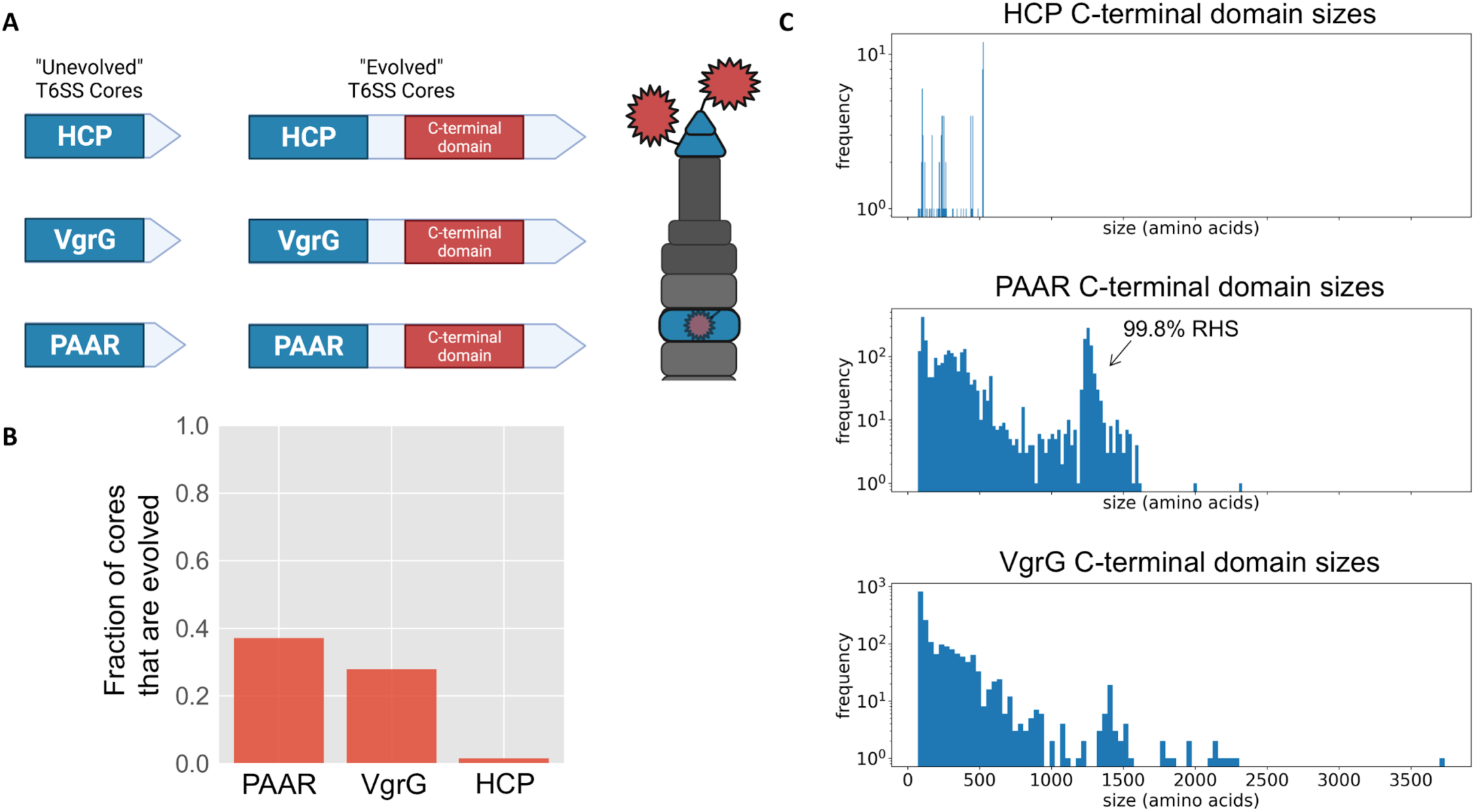
“Evolved” T6SS cores prevalence and sizes. (A) Cartoon of “unevolved” (left) vs. “evolved” T6SS core proteins (middle). “Evolved” cores have a C-terminal domain, which many times is an enzymatic toxic domain, which is inherently loaded onto the T6SS structure, due to the N-terminal core domain (right). (B) Fraction of cores which are “evolved”. (C) Histogram of sizes of C-terminal extensions of HCP, PAAR, and VgrG.

Using our dataset of T6SS cores (Supplementary Data 3), we asked what proportion of cores are “evolved”, i.e. have a C-terminal domain. While it is common for PAAR and VgrG to be “evolved” (38.48% and 26.40% have C-terminal domains, respectively), HCP only quite rarely has “evolved” versions, with only 2.07% of HCP having a C-terminal domain (Figure 3B). We speculate that it may be evolutionarily challenging to fit functional toxic effectors into the small, donut-shaped HCP hexamer for proper packaging, assembly, and delivery. In order to explore whether size constraint may play a role in “evolved” cores, we plotted the sizes of the C-terminal extensions of HCP, VgrG, and PAAR (Figure 3C). HCP has the smallest range of C-terminal domains (between 71 and 528 amino acids, median size 253 amino acids). The vast majority of VgrG C-termini are also less than 500 amino acids (median size 135 amino acids), yet we see some outliers of larger size, up to a maximum of 3663 amino acids. The C-terminal domains of PAAR has a bimodal size distribution, with one group of effector domains less than 500 amino acids, and another large group of 1200-1400 amino acids. This latter group of large effector domains are virtually all annotated as RHS domains, which are large domains that are found in nature to have a variety of C-terminal toxins (Ma, Sun, et al. 2017; Jamet and Nassif 2015) (Figure 3).

We sought to understand how the “evolved” T6SS cores are distributed taxonomically, in order to determine if there is a bias in the distribution of certain “evolved” cores by a given taxonomic group. Strikingly, we can see that “evolved” HCP seem to be overwhelmingly restricted to *Escherichia* (Figure 4A, 10 o’clock to 1 o’clock), which represents 91.4% (1702/1861) of all genomes with “evolved” HCP. This refines our previously mentioned result: “evolved” HCP is scarce, perhaps because it is narrowly distributed taxonomically, and likely appeared only once in an *Escherichia* ancestor. *Salmonella* species seem to have a strong bias towards using mostly “evolved” PAAR (Figure 4A, 1 o’clock to 2 o’clock). Interestingly, *Vibrio* is split into two subpopulations: one that largely uses only “evolved” VgrG, and another subpopulation that only uses “evolved” PAAR (Figure 4A, 4 o’clock to 5 o’clock). The subpopulation that uses only “evolved” VgrG is largely made up of *V. cholerae* (83% of those with “evolved” VgrG only), and the subpopulation that encodes only for “evolved” PAAR is largely made up of *V. parahaemolyticus* (60.78%). We speculate these taxonomic biases may have to do with the fact that these effectors have an N-terminal core domain that needs to be compatible with the underlying T6SS core machinery in order to be loaded and fired. We then asked if it was possible for a genome to encode a T6SS with strictly “evolved” cores, i.e. is it possible to build a T6SS without “regular” cores. We found that virtually no genomes have strictly “evolved” cores (0.4%; 55/11752). Yet, 77% (9087/11752) of genomes contain at least one “evolved” core, showing their extensive utility as part of T6SS function. Looking closer, it is rare to find genomes with strictly “evolved” HCP (0.5%; 63/11289) or strictly “evolved” VgrG (7%; 752/10663), yet it is relatively common to find genomes with all PAAR being “evolved” (27.2%; 2777/10180). This suggests that “evolved” PAAR does not affect efficiency of T6SS assembly, while “evolved” VgrG trimers and “evolved” HCP hexamers may have some cost to T6SS functionality.

**Figure 4.**
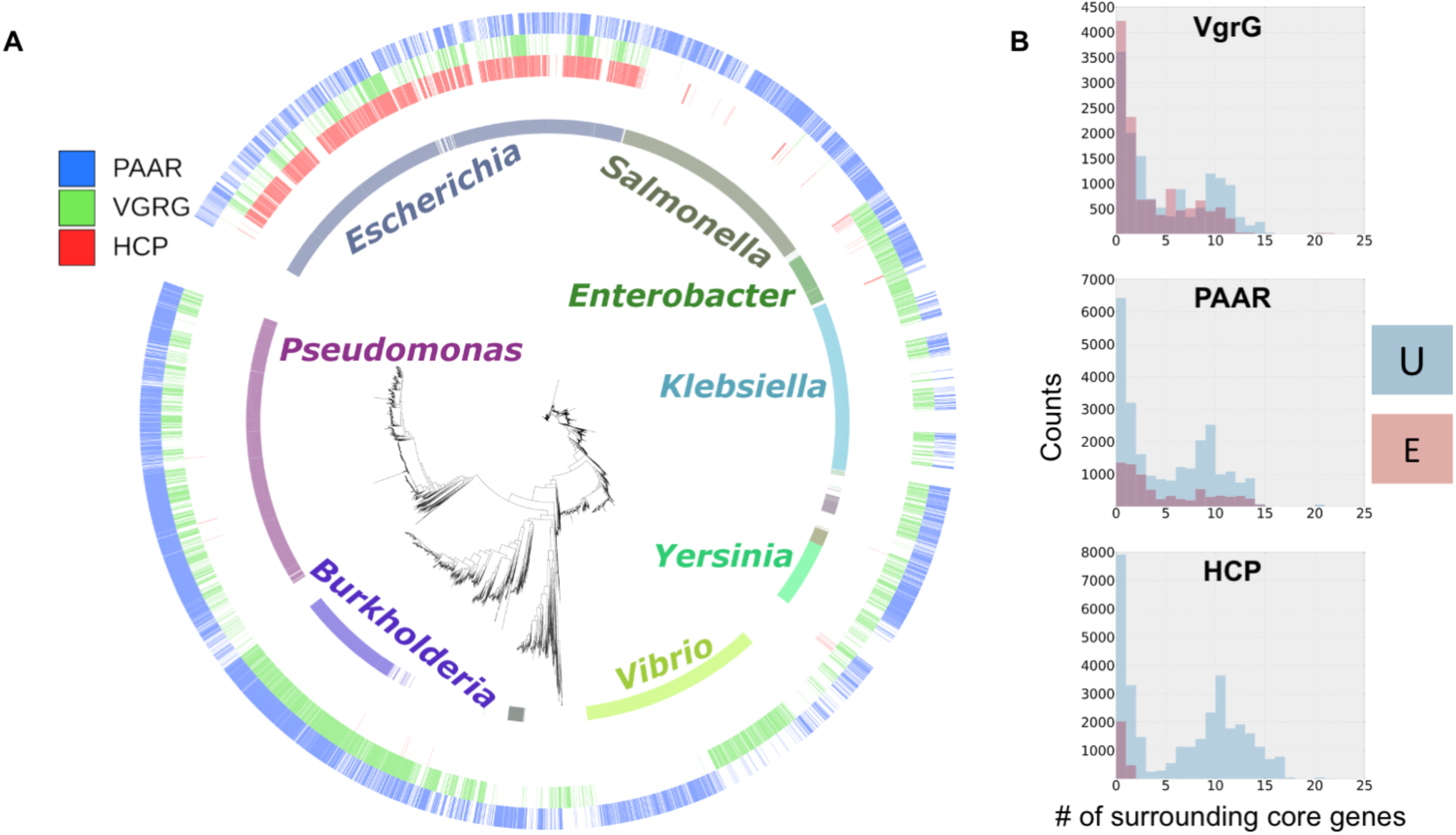
“Evolved” T6SS cores taxonomic and physical distribution. (A) Phylogenetic tree of T6SS+ genomes show how “evolved” PAAR, VgrG, and HCP (outer rings) are distributed across the Genera (inner ring). Only genera with >100 members in the dataset are shown. (B) Histograms of surrounding genes for each core, “unevolved” (red) or “evolved” (blue). Overlapping distributions produce a purple color.

Many times, bacterial genomes contain a main T6SS operon that encodes all the structural genes and effectors, both regular and “evolved” effectors. However, often there is also a genomic “orphan” or “auxilliary” operon, which contains just a small fraction of a T6SS operon, such as an effector gene and its cognate VgrG (Crisan et al. 2019). We asked whether there is a preference for “evolved” T6SS cores to be encoded in the main T6SS operon, or in smaller orphan/auxiliary operons. Regular HCP with no C-terminal domain is encoded both in operons and outside of operons. In contrast, “evolved” HCP have a strong bias for being encoded in orphan operons (98% of “evolved” HCP have 0 or 1 nearby T6SS core gene), with a minority present in main operons (Figure 4B). VgrG and PAAR are found in large numbers in both orphan and main operons, suggesting no bias (Figure 4B).

Since T6Es act generally as antibacterial toxins (Jurėnas and Journet 2021), the identity of the C-terminal domains of “evolved” T6SS cores are of high interest to science and industry. We surveyed all pfam domains encoded in C-terminal domains of HCP, VgrG, and PAAR. Overall, “evolved” HCP has a very narrow variety of known pfam domains, as compared to VgrG and PAAR, which is somewhat expected due to the scarcity of these domains (Figure 5). The most common HCP C-terminal domains encode Colicin- DNAses, as has been seen previously (Ma, Pan, et al. 2017; Howard et al. 2021) (Figure 5). The vast majority of VgrG C-terminal domains have no annotation, likely representing novel toxins. The C-terminal domains of “evolved” PAAR most frequently encode RHS repeats, that are usually encoded upstream of a toxic domain (Jamet and Nassif 2015; Ma, Sun, et al. 2017; Donato et al. 2020). Taken altogether, we conclude that “evolved” HCP has very different characteristics as compared to “evolved” VgrG and PAAR. “Evolved” HCP are very rare overall and appear nearly exclusively in a single genus. When “evolved” HCP do occur, they have comparatively short C-terminal domains, with a narrow variety of known C-terminal pfam domains, likely representing rare HCP-toxin fusion events. Furthermore, HCP are mostly encoded in orphan operons, rather than in main T6SS operons.

**Figure 5.**
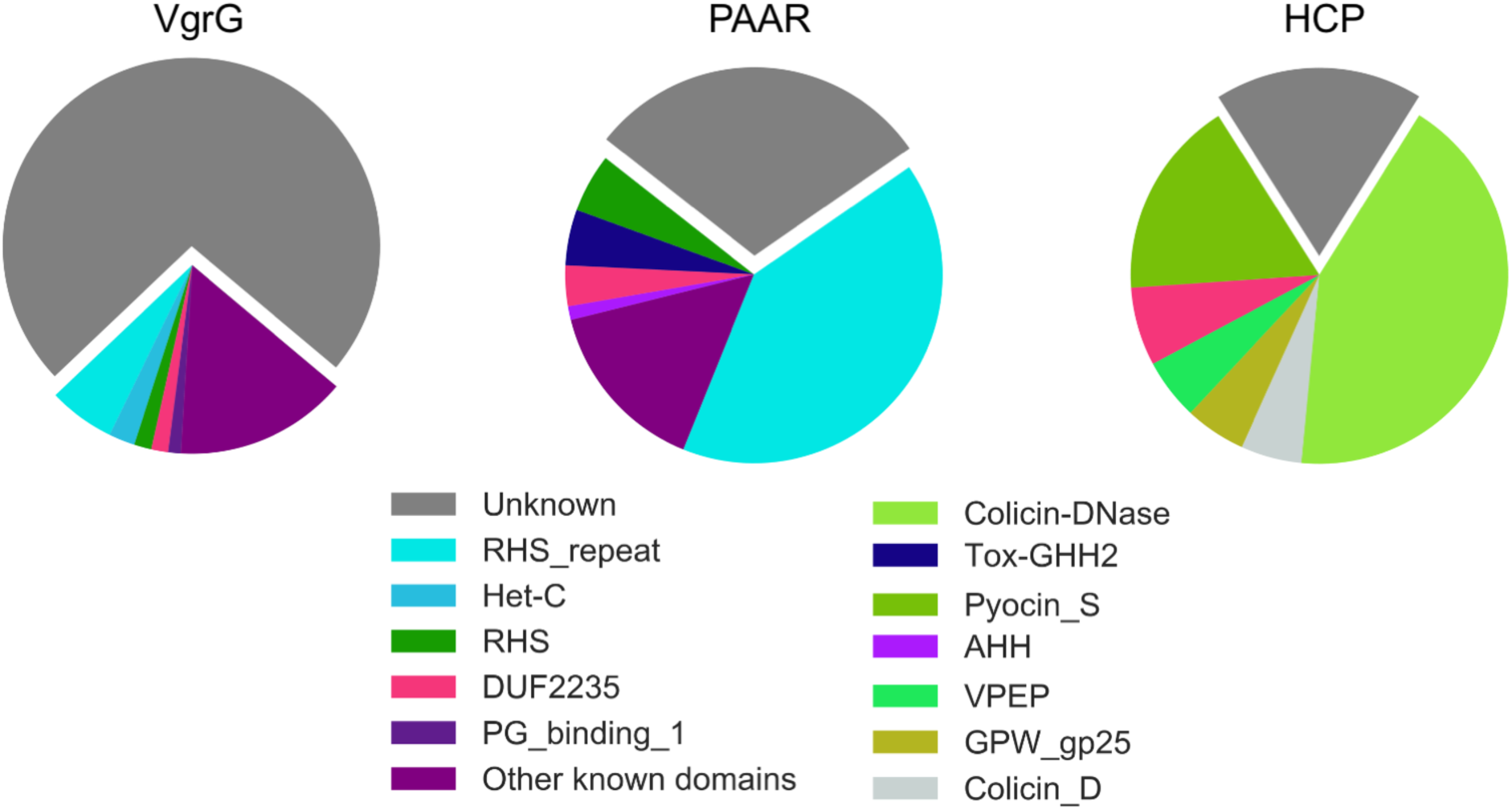
Many C-terminal T6SS core extensions have no annotation. Annotations of C-terminal domains using the pfam database. To avoid phylogenetic bias in plotting proportions of C-terminal domains, one genome per clade (average distances < 0.1) was sampled. This sampling was performed 100 times, each time the makeup of the C-terminal domains of “evolved” cores was noted. After the 100 iterations, the average counts of C-terminal domains were plotted here (average n ≈ 52, 576, and 21 for VgrG, PAAR, and HCP, respectively). Domains with less than 1% occurrence on average were considered below threshold for plotting and grouped together into “Other known domains” (purple). Names in the legend are the pfam database short names.

### Discovery of a novel toxic putative “evolved” HCP - Type VI immunity pair

HCP C-terminal extensions include extensions that are not mapped to known pfam domains (18%; Figure 5, right). This was intriguing and represented an opportunity to discover novel antibacterial T6Es. We designed a pipeline to discover and validate these toxins (Figure 6A): first, we pulled all HCP genes from T6SS-encoding genomes from our database and labeled evolved HCP genes by having at least a 70 amino acid extension after the HCP pfam domain. Then we clustered all the extensions into families and searched for candidate domains that were functionally unannotated, phylogenetically scattered, and encoded next to a cognate immunity protein. We called one gene that fit this profile “Eh1” (“Extended HCP1”) (Supplementary FIgure 1). Eh1 has no annotated function, is present in various genera, such as *Citrobacter* and *Salmonella*, and has synteny with a small gene encoded downstream to it (Supplementary Figure 1, 2). We expressed Eh1 heterologously to test its toxicity to prokaryotic cells. Uninduced, Eh1 is already highly toxic to *E. coli* (Figure 6B, left), suggesting leaky expression is enough to stop bacterial growth. However, upon induction, we see an even higher level of toxicity (Figure 6B, right). Since T6Es are mainly known to target prokaryotic structures that are widely conserved, the bacteria that encode these effectors frequently encodes an immunity protein that neutralizes the T6Es activity. Eh1 had a gene encoded downstream to it, which we hypothesized served as the immunity gene. Indeed, upon co-expression of Eh1 and its putative immunity protein, that we termed Ehi1 (“Extended HCP1 immunity”), we saw a neutralization of Eh1’s toxic activity (Figure 6B), both while the effector is induced and uninduced. Leakiness of the Ehi1 plasmid promoted cell rescue, even without induction by IPTG.

**Figure 6.**
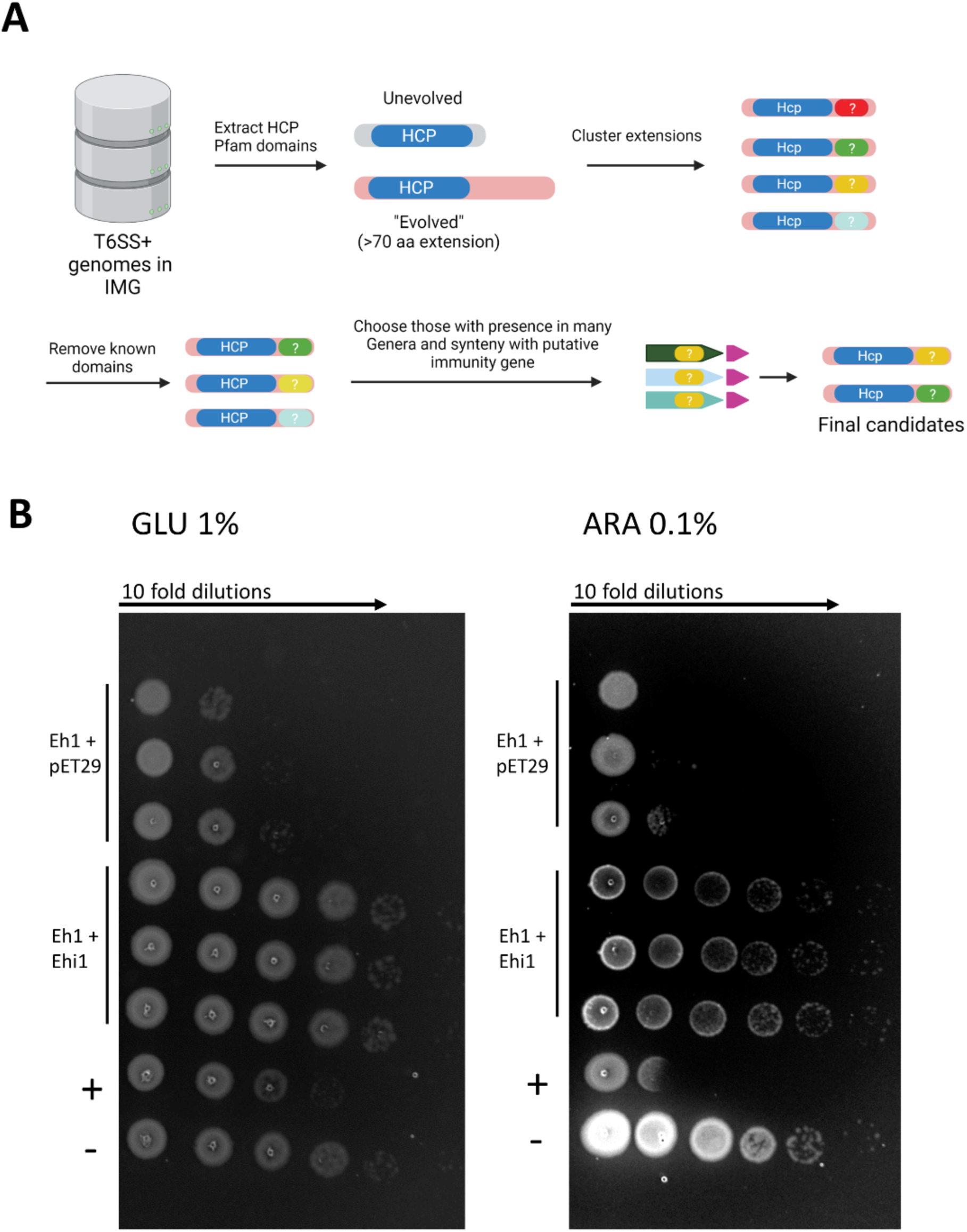
HCP C-terminal domain with unknown annotation is toxic and is neutralized by immunity protein. (A) Pipeline for discovery of novel T6Es in C-termini of HCP. HCP genes from T6SS genomes were considered “evolved” if they had a >70 amino acid extension. To remove redundancy, C-termini were clustered. C-terminal domains with known annotations were discarded, while those with wide phylogenetic presence, and with evidence of a putative immunity protein encoded nearby were kept. Candidate effectors were synthesized and cloned into an inducible vector. (B) Drop assay of *E. coli* heterologously expressing both Eh1 in pBAD24 and Ehi1 in pET29b, or both Eh1 in pBAD24 and an empty pET29b in uninduced (left, 1% Glucose) and induced (right, 0.1% Arabinose) conditions. A known toxin was used as positive control (“+”) and a negative control (empty pBAD24 and pET29b) (“-”) was used to show vector non-toxicity.

### Discovery of nine toxic putative T6Es in an adaptor-based algorithm

The intriguing presence of the DUF4123 domain in the C-termini of “evolved” VgrG inspired us to develop a second algorithm to identify T6Es. DUF4123 does not have a known T6E activity, rather, it is an adaptor of the T6SS (sometimes called a T6SS “chaperone”), which generally helps load the T6SS machinery with effectors (Unterweger et al. 2015; Liang, Moore, and Wilton 2015). Effectors are usually encoded downstream of chaperones, as shown with DUF4123 in Liang et al. (Liang, Moore, and Wilton 2015). We designed a pipeline for discovery of novel adaptors and T6Es using these concepts. Other computational pipelines to discover T6Es have been tried (Russell et al. 2012; Whitney et al. 2013; Salomon et al. 2014; Jana et al. 2019; Jiawei Wang et al. 2018), but they usually rely on sequence similarity to known T6Es or genetic linkage to T6SS core genes. The novel approach that we will describe below is independent of these features.

The goal of the pipeline was to use a computational approach to systematically discover novel adaptors and toxins. Although there has been one study that uses an adaptor-based algorithm, there was no experimental validation of the findings (Liu et al. 2020). We hypothesized there may be more hidden T6SS adaptors encoded in the C-termini of T6SS cores. We marked any predicted C-terminal domain that lacks known pfam annotation as a possible T6SS adaptor (Figure 7A, left). We searched for standalone versions of these adaptor domains. Namely, we looked for occurences of the putative adaptor gene in a single-domain gene in another T6SS-encoding genome (Figure 7A, right). The result of this analysis gives us examples of putative adaptors that sometimes were encoded in orphan loci, and even sometimes in “super orphan” loci, i.e. zero T6SS core genes are in the orphan locus. In other words, these putative adaptors are not genetically linked to any T6SS core gene and we assume that they form non-covalent contacts with T6SS proteins that are encoded elsewhere in the genome.

**Figure 7.**
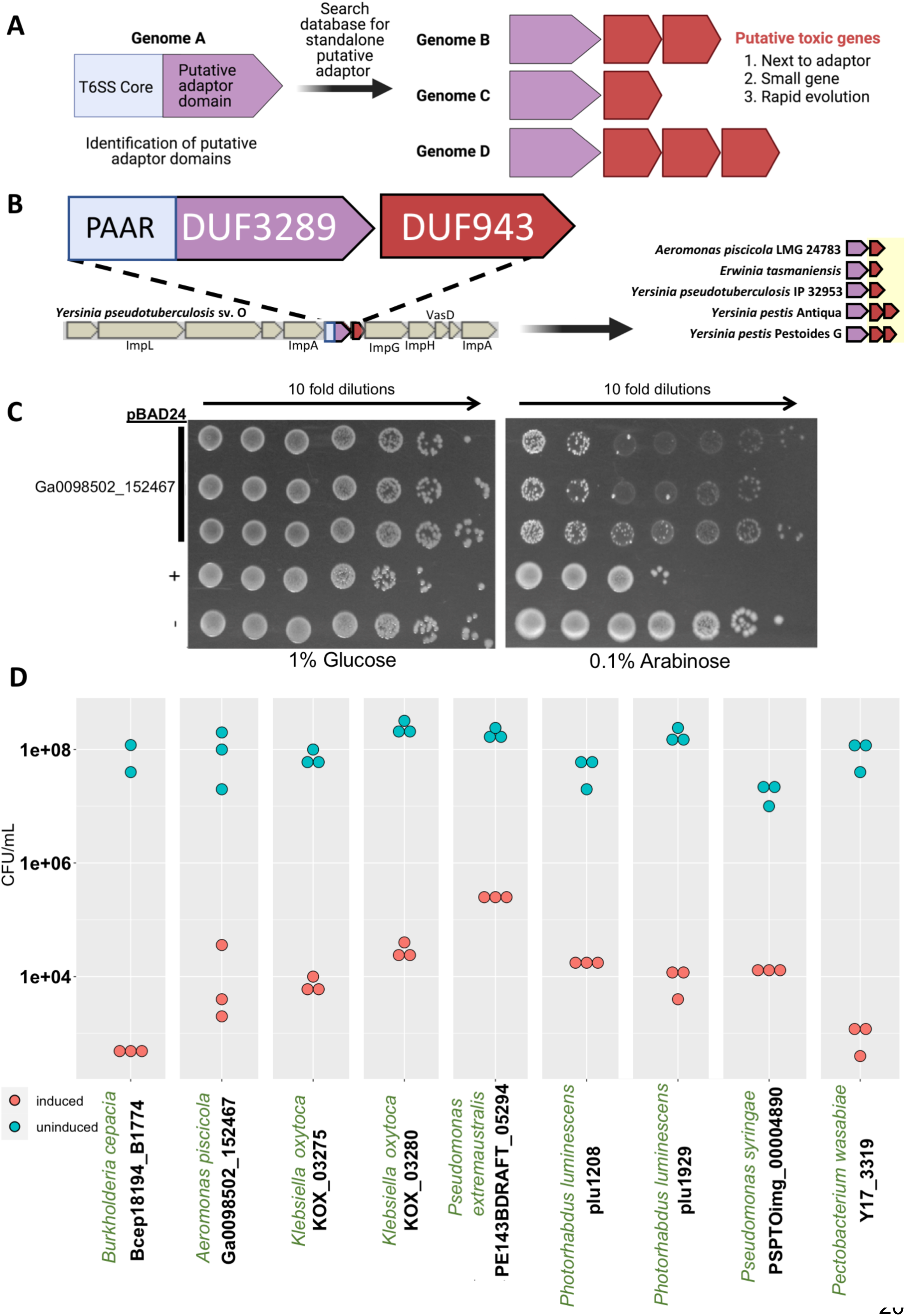
Toxicity of predicted T6Es to *E. coli*. (A) Adaptor- and rapid evolution-based algorithm. Left side shows a protein encoded by a gene in Genome A with an N-terminal T6SS core domain (light blue) and a C-terminal unknown domain, which is considered a putative adaptor domain (purple). Using this C-terminal domain as a query in a sequence database, we searched specifically for standalone versions of this domain, i.e. genes with the putative adaptor domains with no T6SS core domain (right, purple). Encoded next to these adaptors are putative T6Es (red), which have signs of rapid evolution (copy number variation). (B) An example result from the algorithm. In a T6SS operon in *Yersinia pseudotuberculosis* sv. O (left) is a gene that corresponds to a protein with an N-terminal PAAR domain (light blue), and a C-terminal domain with unknown function DUF3289 (purple), which we considered a putative adaptor. Upon looking for orthologs of the DUF3289 domain in other genomes, we identified multiple standalone versions, with a linked gene undergoing rapid evolution (right, red genes). (C) A representative drop assay. Empty pBAD24 plasmids (-), or positive control (a known toxin, +), or putative T6E from *Aeromonas* (see B, right side) were transformed into *E. coli* BL21 (DE3) pLysS. Colonies were grown overnight, and were normalized and serially ten fold diluted and dropped onto agar plates with repressing 1% Glucose (left) and inducing 0.1% Arabinose (right). (D) Quantification of the drop assay, as well as for other putative T6Es which were toxic to *E. coli*. X-axis lists genomes in green and locus tags of the tested genes in black.

This list of standalone putative adaptors was then surveyed for genes that are located adjacent to the putative adaptor. We looked for genes that are small and are rapidly evolving in terms of copy number variations between phylogenetically closely-related genomes (Figure 7A, right). We hypothesized that these genes were putative T6Es because it is known that T6Es are encoded downstream of adaptors (Liang, Moore, and Wilton 2015). We used the rapid evolution filter as a molecular signature of an “arms race” between attacking bacterial T6E genes and the genome of an attacked bacteria that likely evolves resistance against T6Es.

For example, *Azospirillum lipoferum* R1C encodes an evolved PAAR with a C-terminal domain, which is a putative adaptor domain (Supplementary Figure 3). This C-terminal domain exists as a standalone gene (that we previously termed as “Hyde2”) in *Acidovorax avenae subsp. citrulli* AAC00-1 (Supplemental Figure 3). Encoded right next to it is Hyde1, which is known to be highly toxic to *E. coli* in heterologous expression experiments, suggesting it may be a T6E (Levy et al. 2017). On the other hand, Hyde2 is only very slightly toxic to *E. coli*, suggesting it is not a T6E (Levy et al. 2017). Based on the facts that (i) a domain with high similarity to Hyde2 is attached to a core T6SS domain (PAAR), (ii) it is not strongly toxic to *E. coli,* and (iii) a toxic rapidly evolving gene is encoded next to it (Supplemental Figure 3), we conclude that Hyde2 is a putative T6SS adaptor, and that Hyde1 is a putative T6E. Future in depth studies, such as competition assays with mutant strains and co-IP experiments are needed for further validation of this model.

Using this schema, we searched over 11,000 T6SS-encoding genomes, which yielded 43,546 cores with extended C-terminal domains (Supplementary Figure 4). After clustering these C-termini into 2324 unique families, we removed any known annotation to known adaptors, effectors, toxins, and enzymes, leaving 726 families with no functional annotation. We considered these possible adaptor sequences, while keeping in mind they may be effector domains. As explained above, we then searched for standalone gene versions of these putative adaptors, specifically for those encoded in orphan and “super orphan” loci, leaving 89 possible candidates. We then manually screened candidates and then evaluated each locus for presence of a small gene with signatures of rapid evolution; these were labeled as putative T6Es. We defined the rapid evolution by copy number variation of the genes between related genomes. Supplementary Figures 5-12 detail some of our candidates. One example of the output of this pipeline is shown in Figure 7B, which shows a particular gene pair in a T6SS operon in the pathogen *Yersinia pseudotuberculosis* sv. O. The upstream gene has an N-terminal PAAR domain, and attached to it is a C-terminal domain with pfam annotation DUF3289, which has no known function (Figure 7B). We hypothesized that DUF3289 is a putative adaptor. Encoded next to this “evolved” PAAR is a gene which encodes a protein with a DUF943 domain, also with no known function (Figure 7B). Upon searching other bacterial genomes for orthologs of the DUF3289 domain, we found that in multiple genomes, there exists a standalone gene version, containing a DUF3289 domain only. These standalone versions of the DUF3289-encoding putative adaptors also had synteny with DUF943-encoding genes (Figure 7B, right). Furthermore, we observed a copy number variation between similar loci, leading to our hypothesis that DUF943-containing proteins are putative T6Es. One of the standalone DUF3289-DUF943 pairs was synthesized into inducible expression vectors (Figure 7B, genes from *Aeromonas piscicola* LMG 24783), along with various other candidates (Table 1, Supplementary Table 2). Heterologous expression assays in *E. coli* were then used to rapidly validate our computational predictions. Upon expression of the DUF943-encoding putative effector Ga0098502_152467 from *Aeromonas piscicola* LMG 24783 in *E. coli,* we observed strong antibacterial activity (Figure 7C). The adjacent DUF3289 encoding gene, the putative adaptor, was not toxic to *E. coli* (Supplementary Table 2). Overall, we screened 20 candidates, and our results overall showed seven examples of non-toxic putative adaptors with downstream T6Es (Supplementary Table 2), and 9 putative T6Es that had a toxic effect on *E. coli* (Figure 7D, Table 1, Supplementary Figures 13-20). Interestingly, 5 of the candidate T6Es that we experimentally validated have a predicted transmembrane domain (Table 1). Protein modeling reveals highly hydrophobic helices in the structures (Supplementary Figure 21). Therefore, we speculate that these proteins are killing microbes as they degrade the bacterial membrane or destroy the proton motive force by forming pores. A transmembrane domain is also present at the N terminus of Hyde1 putative T6E (Levy et al. 2017).

**Table 1.**
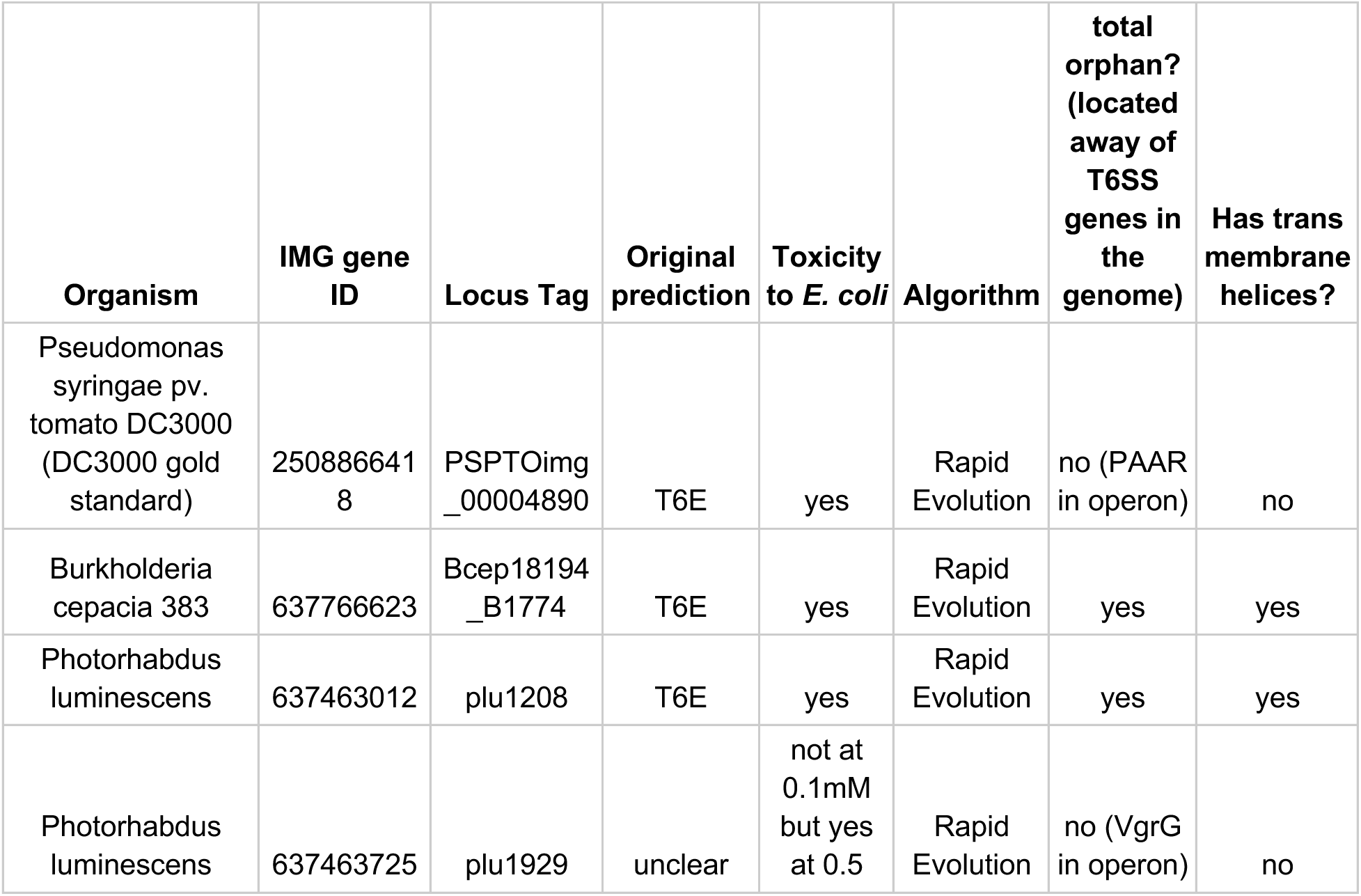
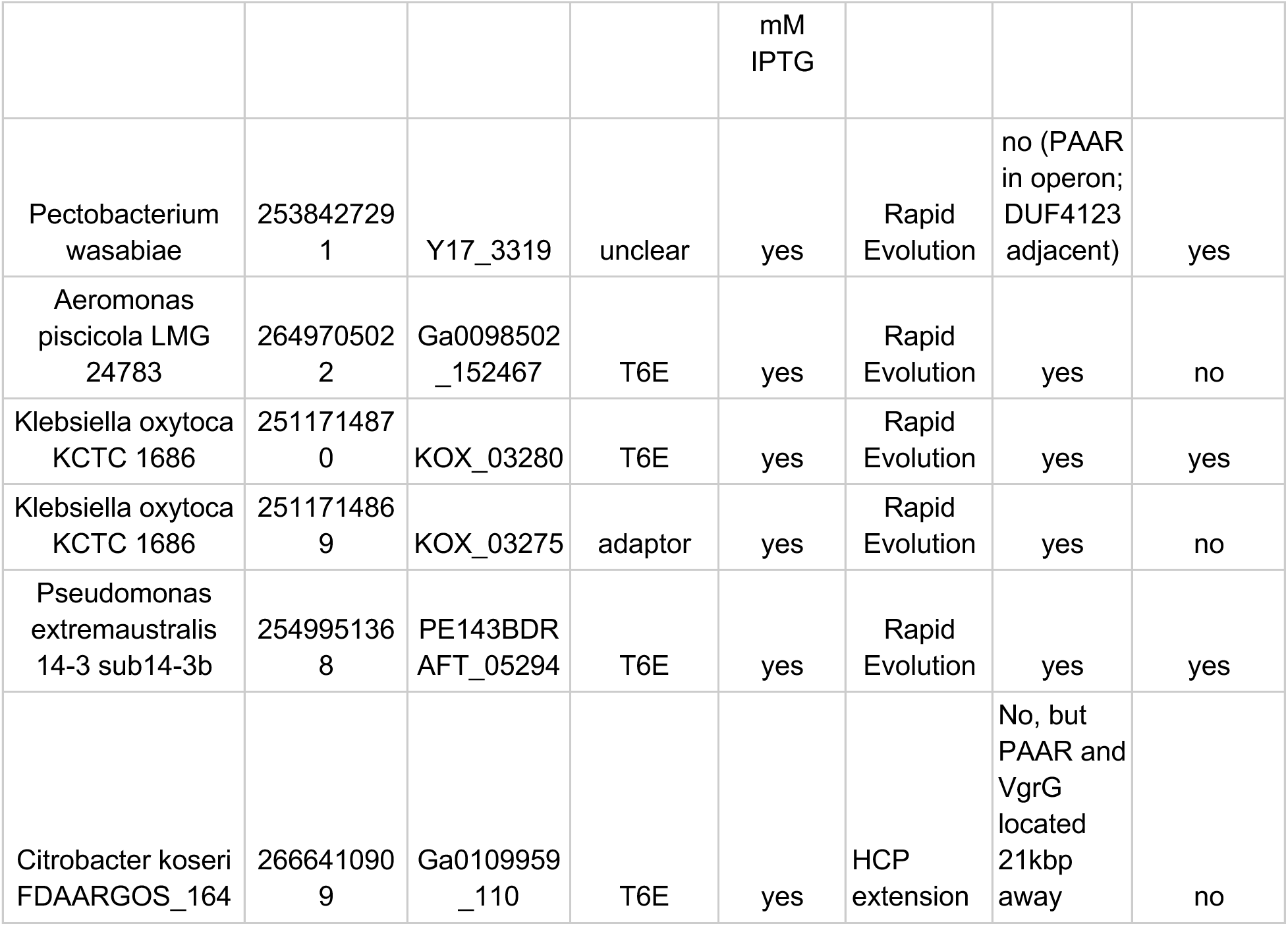
Genes toxic to E. coli are defined as putative T6Es.

Although these experiments are only a first indication of possible T6Es, we underscore the fact that a number of species we studied have no known T6SS effector, as per a search in the SecReT6 database (J. Li et al. 2015). This is important because many of these species have important effects on medicine and economics. For example, *Aeromonas piscicola* is found in diseased fish, *Klebsiella oxytoca* is a drug resistant opportunistic pathogen, *Burkholderia cepacia* is an opportunistic human pathogen, *Pectobacterium wasabiae* is a plant pathogen, and *Citrobacter koseri* is a known opportunistic human pathogen (Beaz-Hidalgo et al. 2009; Darby et al. 2014; Singh, Cariappa, and Kaur 2016; Mahenthiralingam, Urban, and Goldberg 2005; Kim et al. 2009; Deveci and Coban 2014). First hints of a possible T6E in these species allows for other laboratories to quickly study these potentially important virulence factors that may affect colonization and/or pathogenesis.

Taken altogether, we have used the birds-eye-view analysis of “evolved” T6SS cores to design two algorithms for discovery of putative novel T6Es. The latter algorithm is sequence and location independent, and can identify “super orphan” loci of putative T6Es. By using bioinformatic analysis, we were able to design a high throughput approach that led to candidate T6E discovery in eight organisms. Using heterologous expression experiments we showed toxicity in 10 genes, which we label as putative T6Es. This is in contrast to deep, yet narrow studies of T6Es which often result in discovery of a single T6E family from one species. Our approach is broader and may provide a faster way to screen for genes for further basic and applied research.

## Discussion

In this study, we used a combined bioinformatic-experimental approach to screen and identify 10 putative T6Es. Not only did we survey the universe of C-terminal extensions of “evolved” T6SS cores, we used the information to develop two computational approaches that identify novel T6SS effectors in thousands of bacterial genomes. Notably, using the second algorithm, the adaptor-based algorithm, we were able to detect some examples of effectors independent of sequence homology to known effectors and independent of proximity to T6SS cores. We tested the toxicity of several candidate effectors that we predicted via heterologous expression experiments in *E. coli* and showed that many are indeed toxic and for one, we have a cognate immunity gene that protects from toxicity. These experiments are common in the T6SS research field to confirm antibacterial or anti-eukaryotic activity. We are aware that in order to define a gene as a *bona fide* T6SS effector we need to provide and evidence of T6SS-dependent secretion, and that the presence of the gene in the attacking strain is correlated with higher fitness in competition assays against a prey strain that lacks the immunity protein. In our analysis, we provided strong genetic evidence that the novel toxins are associated with T6SS, but we did not construct mutants in the encoding strains nor identify the conditions in which the toxins are being secreted. Therefore, to be cautious, we termed the new genes “candidate” or “putative” T6SS effectors. Future experiments are required to confirm that the candidate effectors are secreted in a T6SS-dependent manner and serve for intercellular competition purposes. Nevertheless, using computational evidence and a simple bioassay, we were able to screen and discover a “short list” of putative T6Es for future studies. This process can theoretically be scaled up to characterize the toxicity of all families of unknown C-terminal domains of “evolved” cores, providing an exhaustive list of promising hits for deeper study.

Unsurprisingly, some of the predicted adaptors were toxic in the heterologous expression assay, as it is possible that the C-terminal domains attached to T6SS cores are toxins. Indeed, we also took advantage of this concept for the first algorithm described in this paper (Figure 6A). We therefore expect the downstream putative “effector” of a toxic “adaptor” to possibly be an immunity gene.

We also saw cases where neither the putative adaptor or the putative T6E are toxic, which may be because they are not expressed in the correct compartment, i.e. one or the other may need periplasmic expression. We kept in mind that *E. coli* is likely not the natural target of a specific T6E, so this may also be a reason for predicted effectors having no toxic activity. Nevertheless, non-toxicity of some of the predicted genes demonstrates that the toxicity to *E. coli* is not simply due to overexpression of foreign protein.

One specific putative T6E (plu1929) was not toxic in heterologous expression assays at 0.1mM of IPTG induction. However, upon induction at 0.5 mM, it was indeed toxic (Supplementary figure X, Table 1). We counted this as a possible T6E, as we saw that even at 0.5 mM IPTG its upstream putative adaptor (plu1928) was not toxic, demonstrating that at this higher level of induction, there was still not a general toxicity caused by overexpression (Supplementary Figure 15). We reasoned that plu1929 should be investigated further for evidence of T6SS effector activity, and therefore counted it in our list of putative T6Es (Table 1).

We also noted that for many of the putative effectors, we could not predict cognate immunity genes. This could be due to the fact that the immunity protein was either encoded somewhere else in the genome, but also could be due to non-immunity-protein-based immunity, i.e. by general mechanisms of immunity (Le et al. 2020; Toska, Ho, and Mekalanos 2018; Hersch et al. 2020; Hersch, Manera, and Dong 2020; Flaugnatti et al. 2021).

Our previous study described the extracellular contractile injection system (eCIS) which is enriched in understudied bacterial genera (Alexander Martin Geller et al. 2021; L. Chen et al. 2019). eCIS is structurally and evolutionarily related to the T6SS, with the first being extracellular and the second being membrane-bound. It was also noted that eCIS and T6SS are not frequently encoded together, suggesting an overlapping function (L. Chen et al. 2019). As expected, we saw that T6SS-encoding genomes had an enrichment of well-studied Genera, the opposite of the result of eCIS. We saw the genera that encoded eCIS and for T6SS were largely non-overlapping, yet we saw exceptions. *Dickeya* and *Xenorhabdus* genomes are statistically enriched in both eCIS and T6SS, and *Shigella* genomes are statistically depleted in both eCIS and T6SS (Figure 2) (Alexander Martin Geller et al. 2021; L. Chen et al. 2019). eCIS-encoding bacteria are mostly found in terrestrial and aquatic environments, and are statistically underrepresented in human isolates, and specifically in human pathogens (A. M. Geller et al. 2020; Alexander Martin Geller et al. 2021). Here, we saw the T6SS-encoding genomes are the opposite, with enriched isolation from humans, as well as enriched in a pathogenic lifestyle. Overall, our analysis fits well with previous knowledge about eCIS and T6SS (Alexander Martin Geller et al. 2021; L. Chen et al. 2019). eCIS effectors seem to be loaded in the lumen of the contractile injection system (Desfosses et al. 2019; Ericson et al. 2019). These effector proteins can be quite large, e.g. AFP18 is 2,366 amino acids in length (Hurst Mark R. H., Glare Travis R., and Jackson Trevor A. 2004); Mif1 is 943 amino acids (Ericson et al. 2019). There does not seem to be a size constraint for effector loading into the lumen of the eCIS. However, our current analysis of “evolved” HCP suggests that there is a strict size limit, as well as a noted lack of variety of C-terminal domains as compared to the other T6SS cores. It is possible that “cargo” (non-covalently bound) effectors may have a larger size limit. The large size of eCIS effectors suggests they are unfolded in the lumen of the particle, and that T6SS may not allow for unfolded effectors in the lumen, based on the fact that, in at least one case, large HCP cargo effectors block T6SS-mediated toxicity (Howard et al. 2021). This is sensical if “evolved” HCP are loaded in the process of extension of regular HCP hexamer units into a tube, while eCIS may be loaded with an effector after assembly, although this is currently understudied. It is possible that a study of T6SS subtype iv may provide examples of larger HCP-related T6Es, as it may assemble more like eCIS. Further studies of T6SS assembly with “evolved” HCP are needed.

Our study used T6SS domains TssJ, TssL, and TssM to define the T6SS genomes (Supplementary Table 1). We specifically saw that we captured the most common T6SS subtype i (and a handful of possible subtype ii), but did not include subtypes iii or iv. This is due to the fact that the conserved domains were likely defined mostly on subtype i T6SS. Nevertheless, there have been few general bioinformatic studies of the T6SS since the discovery of this system, and we were able to update the overall picture using 11,000+ genomes. Future studies are needed to properly define conserved domains that belong to subtype iii and subtype iv, making it possible to research their characteristics overall. Even so, there are still large numbers of undiscovered effectors (LaCourse et al. 2018), as we have shown here by providing ten more putative T6Es.

## Materials and Methods

Selected scripts used in search and analyses is available at https://github.com/alexlevylab/T6E_discovery

### Phylogenetic tree construction

Using the IMG database of publicly available bacterial genomes (Markowitz et al. 2012; I.-M. A. Chen et al. 2017, 2019), we searched for genomes with T6SS marker domains TssJ, TssL, and TssM (Supplementary Table 1). Because of use of these marker genes, we expected to mainly find T6SS of subtype i and ii, as TssJ, L, and M are present in those subtypes (Russell et al. 2014).

Phylogenetic trees were constructed using a subset of universal marker genes (Puigbò, Wolf, and Koonin 2009), i.e. 29 COGs (Galperin et al. 2021) corresponding to ribosomal proteins out of 102 COGs in Puigbò et al. (Puigbò, Wolf, and Koonin 2009). To preserve the quality of the tree, genomes missing more than one COG were dropped from analysis. COGs from each organism were aligned separately using clustal omega version 1.2.4 (Sievers et al. 2011) with default settings. Alignments for each COG were then trimmed using TrimAl version 1.4 (Capella-Gutiérrez, Silla-Martínez, and Gabaldón 2009) using the “automated1” flag. Genomes with missing COG sequences were added as gaps into the alignments. Each trimmed alignment was then concatenated by genome, to create an amalgamated 30 COG sequence. This was inputted into VeryFastTree, version 3.0.1 (Piñeiro, Abuín, and Pichel 2020) using standard settings. Trees were visualized on iTOL (Letunic and Bork 2019) and using ggtree and ggtreeextra packages (Yu et al. 2017; Xu et al. 2021). Representative members of each taxonomic Family were used to build the phylogenetic tree in Figure 1, while Figure 4 used all members of T6SS-encoding genomes.

### Enrichment analyses

The taxonomic data from the T6SS database was compared to the number of genera in publicly available genomes of Bacteria from IMG (Markowitz et al. 2012; I.-M. A. Chen et al. 2019). A Fisher exact test was calculated for each Genus, which returned an odds ratio and an associated p-value, the latter of which was corrected using the Benjamini Hochberg method (FDR) for multiple hypothesis testing.

To correct for taxonomic bias that affects enrichment analysis, a population-aware enrichment analysis based on Scoary (Brynildsrud et al. 2016) version 1.6.16 was used. The inputs to Scoary were (1) the guide tree, (2) a presence-absence file, indicating if a genome encodes for T6SS, (3) metadata files that indicate for each genome if they possess a certain characteristic, e.g. whether it was isolated from soil. One guide tree was created per metadata category, based on universal marker genes (see section on phylogenetic tree construction) and only those entries with at least 25 instances in the database were included.

### Search for “evolved” effectors

An “evolved” effector was defined as an HCP, VgrG or PAAR that has an extension of at least 70 amino acids downstream to the core domain (i.e. the pfam domain that defines the HCP, VgrG or PAAR). In the case of “evolved” HCP and PAAR, there are either one or two pfam domains that define the core domain, and all were used to search for “evolved” cores (HCP: pfam05638, PAAR: pfam05488 and pfam13665; Supplementary Table 1). However, VgrG has multiple domains that define it (pfam05954, pfam06715, pfam10106, pfam13296, pfam04717; Supplementary Table 1), so the “core domain” boundaries are less clear. Many times, the conserved domains for VgrG in the COG and TIGRFAM (Haft et al. 2013) domains were found in a given gene, however the pfam domain was not found in that same gene (TIGR domains: TIGR01646, TIGR03361; COG domains: COG3501, COG4253, COG4540, COG3500, COG4379). Because of these issues, we therefore defined the “evolved” VgrG based on length. Using a Gaussian Mixture Model, we searched for two distributions in all VgrG based on amino acid length, “unevolved” and “evolved”. Since most of the “evolved” VgrGs having a known fused C-terminal domain were quite large, we identified a threshold between the two distributions. We used the Scikit-learn library (Pedregosa et al. 2011) with the function GaussianMixture with parameters of “full” covariance. In order to define the threshold for the “evolved” VgrG, we took the mean length of the “unevolved” VgrG distribution (724aa) and added one standard deviation (72aa), which summed to 796aa length. In other words, all genes with any VgrG pfam domain that is greater than 796 amino acids in length were considered “evolved”.

### Counting “evolved” percentage

In order to see which percentage of each core is “evolved” without phylogenetic bias, we collapsed the phylogenetic tree into groups of leaves with <0.1 branch lengths and sampled one taxon each iteration from each sub-group. Then, the percentage of “evolved” cores was counted. We sampled for 10,000 iterations, and took the final average number of “evolved” cores for each core.

### “Evolved” HCP effector algorithm

All HCP genes were extracted from 11,310 T6SS-encoding genomes, based on the HCP pfam domain (pfam05386). “Evolved” HCPs were defined as described above. The start coordinate of C-terminal extensions of HCP was defined based on the end coordinate of the alignment of the HCP pfam domain. C-terminal extensions with known pfam annotations were filtered out, and all unknown C-terminal extensions were clustered using CD-HIT (W. Li and Godzik 2006; Fu et al. 2012) version 4.8.1 with at least 60% identity and at least 70% coverage of both representative and query (n=155 clusters). Cluster representatives were then searched in HHpred (Söding, Biegert, and Lupas 2005). Representatives that had a hit for a known effector in HHpred were removed. The remaining representatives with unknown function were queried using BLAST (Camacho et al. 2009) against the nr database with default parameters. Queries that had subjects in several Genera (Supplementary Figure 2), and had putative immunity genes (small downstream genes with synteny to putative effectors) were chosen as candidates and were synthesized.

### Adaptor-based “super orphan” algorithm

Since this algorithm depends on detection of rapid evolution between phylogenetically related genomes, the search space for this pipeline began with genomes that were clustered into similarity groups (by 95% Average Nucleotide Identity), n = 43,136 genomes. We used genome clusters which had at least 50% of the genomes with the full complement of marker genes TssJ, TssL, TssM (Supplementary Table 1). This resulted in 11,310 T6SS-encoding genomes.

From these genomes, HCP, VgrG, and PAAR genes were extracted (n = 148,414). Those with C-terminal extensions greater than 100 amino acids and less than 500 amino acids were taken for further analysis (this is because most C-terminal extensions were in this range, and large outliers were not our focus in this study). The boundaries were defined by the start and end alignments of the pfam HMM profiles with each gene. This resulted in 43,546 HCP, VgrG, and PAAR that were considered “evolved”. We then clustered the C-terminal domains using CD-HIT version version 4.8.1 using parameters for >=65% identity and >= 80% coverage on both query and subject. We then submitted the representative sequences of the C-termini to the NCBI CDD (Marchler-Bauer et al. 2015) search, which identifies conserved domains. We were able to label C-termini as having a known toxic enzymatic domain, or those without a known function. Those with known enzymatic/toxin domains were discarded. This left 726 clusters with putative T6SS adaptor function (or toxic effector function). Using DIAMOND (Buchfink, Xie, and Huson 2015; Buchfink, Reuter, and Drost 2021) version 2.0.4.142, we searched the IMG database using the “blastp” subcommand with default parameters for standalone versions of these C-termini, i.e. genes with the putative adaptor domain only (not any core domain). C-termini with no standalone domain were dropped from analysis, as we wanted to design an algorithm that could identify effectors and adaptors in “super orphan” loci. A “super orphan locus” is a T6SS related genes that are located away of any known T6SS gene, in comparison to an orphan operon which may have a few T6SS genes, but not the full T6SS operon. 89 adaptor clusters were found to be encoded in orphan or super orphan operons. Sixteen of these adaptor clusters were manually chosen for downstream analysis. Of these, seven were found to have copy number variation. This was measured by taking genome clusters, and aligning homologous regions, and looking for signs of variation in copy number.

### Protein modeling and visualization

Amino acid sequences of the T6Es were submitted to the webserver of Robetta, using the RoseTTAfold option (https://robetta.bakerlab.org/submit.php) (Baek et al. 2021).

### Drop assays

Putative T6Es were synthesized (codon optimized for *E. coli*), and cloned into either pET28, pET29, or pBAD24 plasmids by Twist Bioscience. The plasmids were then transformed into *E. coli* BL21 (DE3) pLysS strain using either heat shock or electroporation. Overnight cultures of the strains harboring the vectors of interest were grown in LB containing the proper selection: Kan or Amp for pET28/29 or pBAD24, respectively. The cultures were normalized to 0.5 OD600 and subsequently serially ten-fold diluted. Dilutions were spotted on LB agar containing the proper selection and inducer (100 μM IPTG; 0.1% Arabinose) or repressor (1% glucose) and the plates were incubated overnight at 37 degrees Celsius.

For testing of immunity genes, the plasmids of interest, one either pET28 or 29, and the other pBAD24, were co-transformed into *E. coli* BL21 (DE3) pLysS using electroporation. The doubly-resistant colonies were then grown overnight and diluted and plated as mentioned above, except that plates had either 1% Glucose, 1% Glucose and 0.1 mM IPTG, 0.1% Arabinose, or 0.1% Arabinose and 0.1 mM IPTG.

### Visualization

Many figures (3A, 6A, and 7A) were produced using BioRender (BioRender.com).

### Supplementary Information

#### Data

**Supplementary Data 1. Genomes with T6SS marker genes.** Attached supplementary data file with accession IDs and phylogeny of the 11,832 genomes with T6SS marker genes.

**Supplementary Data 2. Scoary enrichment/depletion analysis data.** The data corresponding to Figure 2. The highlighted columns were plotted. The outputs are described in the manual for scoary (https://github.com/AdmiralenOla/Scoary#output)

**Supplementary Data 3. T6SS cores gene data.** IMG gene IDs for core genes and their associated pfam domains, both core and C-terminal domains. Number of genes of T6SS genes surrounding each gene is listed. Gene architecture shows which domains make up the gene from N- to C- terminus.

## Acknowledgements

The authors thank Noam Dotan for help with phylogenetic trees and scripting. AL is generously supported by the Israeli Science Foundation (Grants #1535/20, #3300/20), Alon Fellowship of the Israeli council of higher education, The Hebrew University - University of Illinois Urbana-Champaign seed grant, the Israeli Ministry of Agriculture (Grant 12-12-0002), and ICA in Israel. AMG is generously supported by the Kaete Klausner Scholarship and was supported by a scholarship from the Israeli Ministry of Aliyah and Integration.

---

**Supplementary Figure 1.**
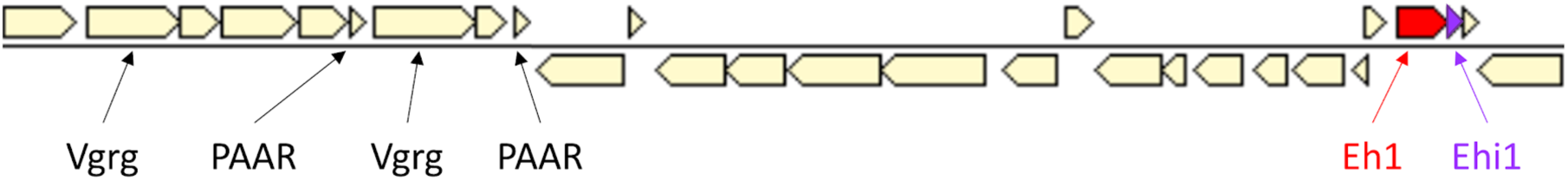
Eh1 and Ehi1 are not part of a T6SS operon, although core genes are nearby. *Citrobacter koseri* FDAARGOS_164 (IMG taxon ID: 2663763285). Eh1 coordinates: 1329871 - 1330971 (locus tag: Ga0109959_1101377). Ehi1 coordinates: 1330955 – 1331209, (locus tag: Ga0109959_1101378).

**Supplementary Figure 2.**
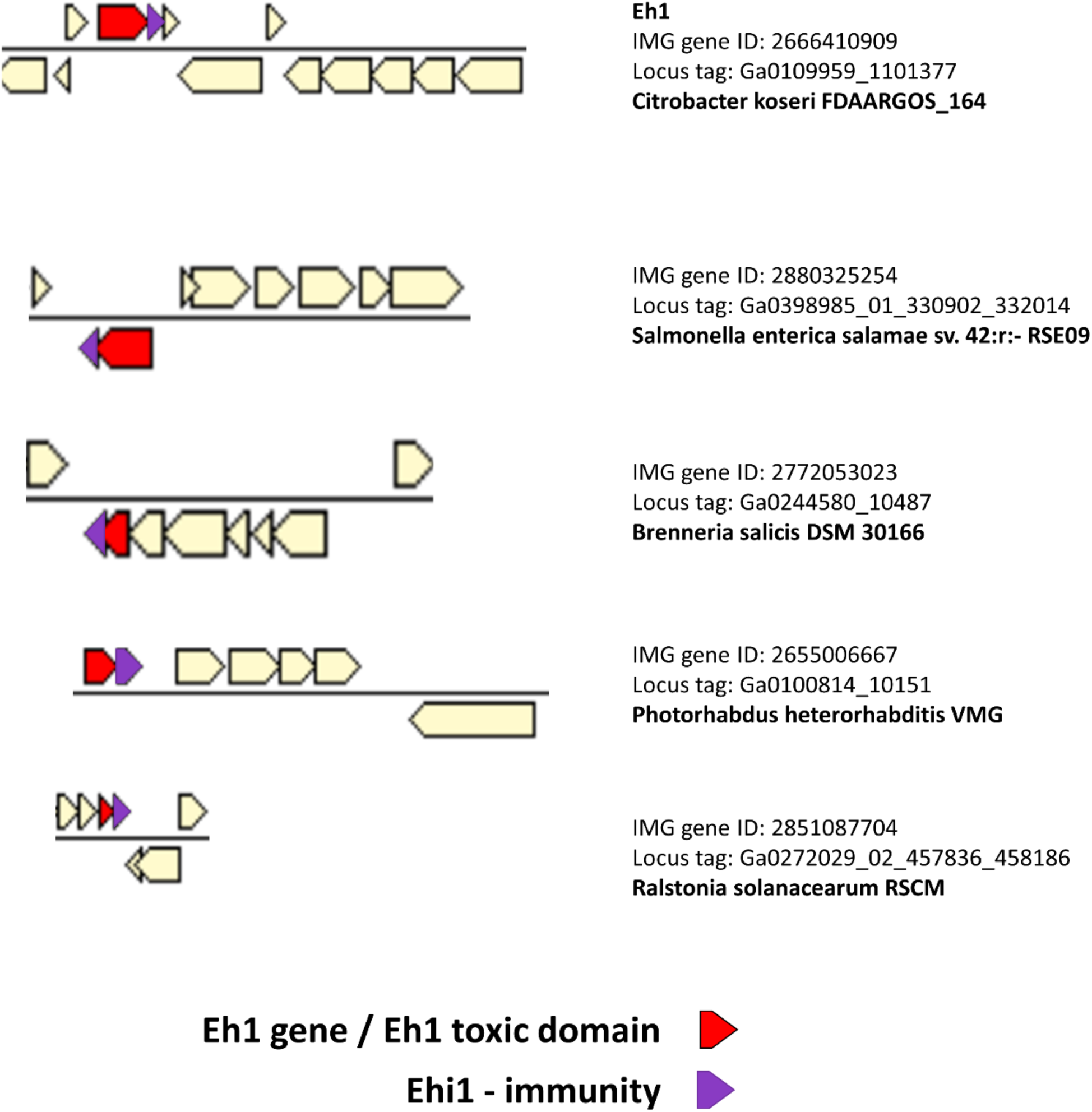
BLAST hits of Eh1 candidate extension exhibit synteny. Eh1 effector domain BLAST hits were found in various Genera, and had synteny with a downstream putative cognate immunity gene. Bottom three are standalone versions, i.e. effector domains only, without an N-terminal HCP.

**Supplementary Figure 3.**
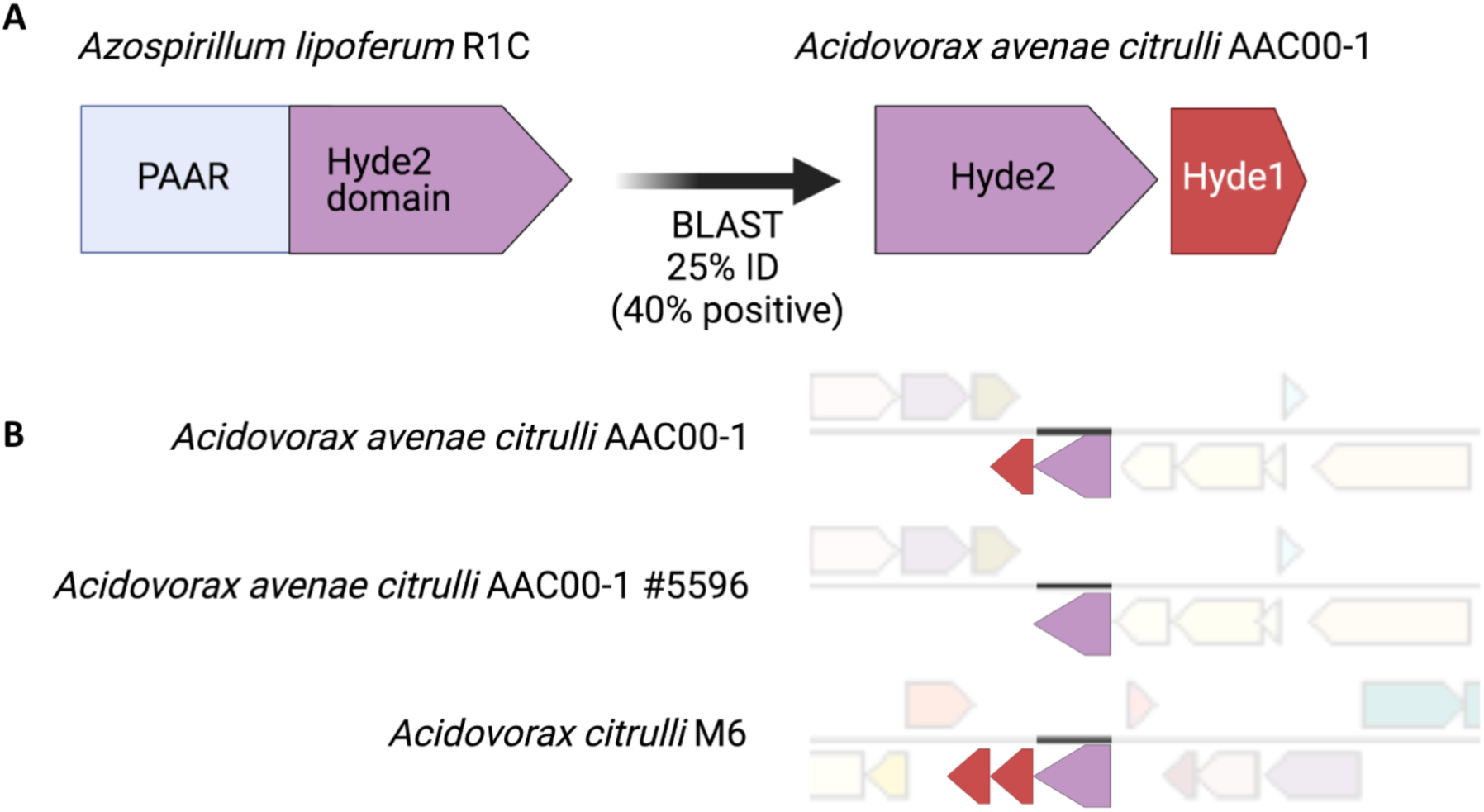
Small genes with signatures of rapid evolution encoded next to T6SS adaptors are putative effector toxins. (A) An example of a C-terminal core domain, Hyde2 (pale blue), connected to an N-terminal PAAR domain (purple), which has a standalone version next to small gene, Hyde1, that is toxic to *E. coli* (Levy et al. 2017). (B) Three loci from closely related species that encode Hyde2 orthologs (purple) and their associated Hyde1 orthologs (red) display copy number variation, a sign of rapid evolution.

**Supplementary Figure 4.**
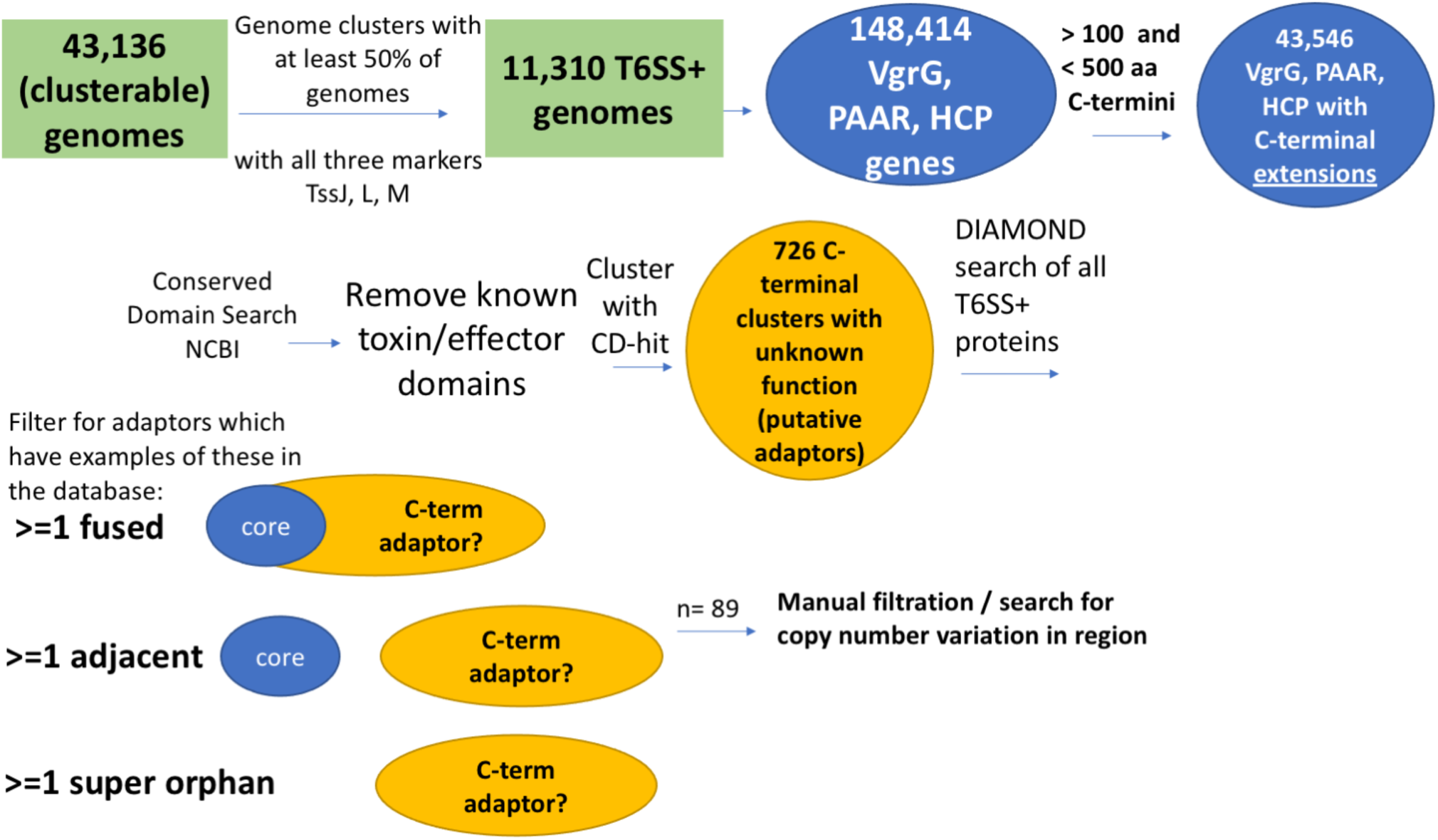
Rapid evolution algorithm. Genomes with known pairwise ANI values were filtered for the presence of T6SS marker domains TssJ, L, and M. In those genomes, genes with marker domains for core T6SS genomes were identified, and filtered for those with extensions >100 amino acids and <500 amino acids. C-terminal domains with annotations for known toxic/effector domains were removed. C-termini were clustered and representatives were used for search for standalone versions of the putative adaptors in other genomes. Those with orphan and super orphan forms were chosen and screened for copy number variation. Selected candidates were synthesized and cloned into inducible vectors and tested for toxicity by heterologous expression.

**Supplementary Figure 5.**
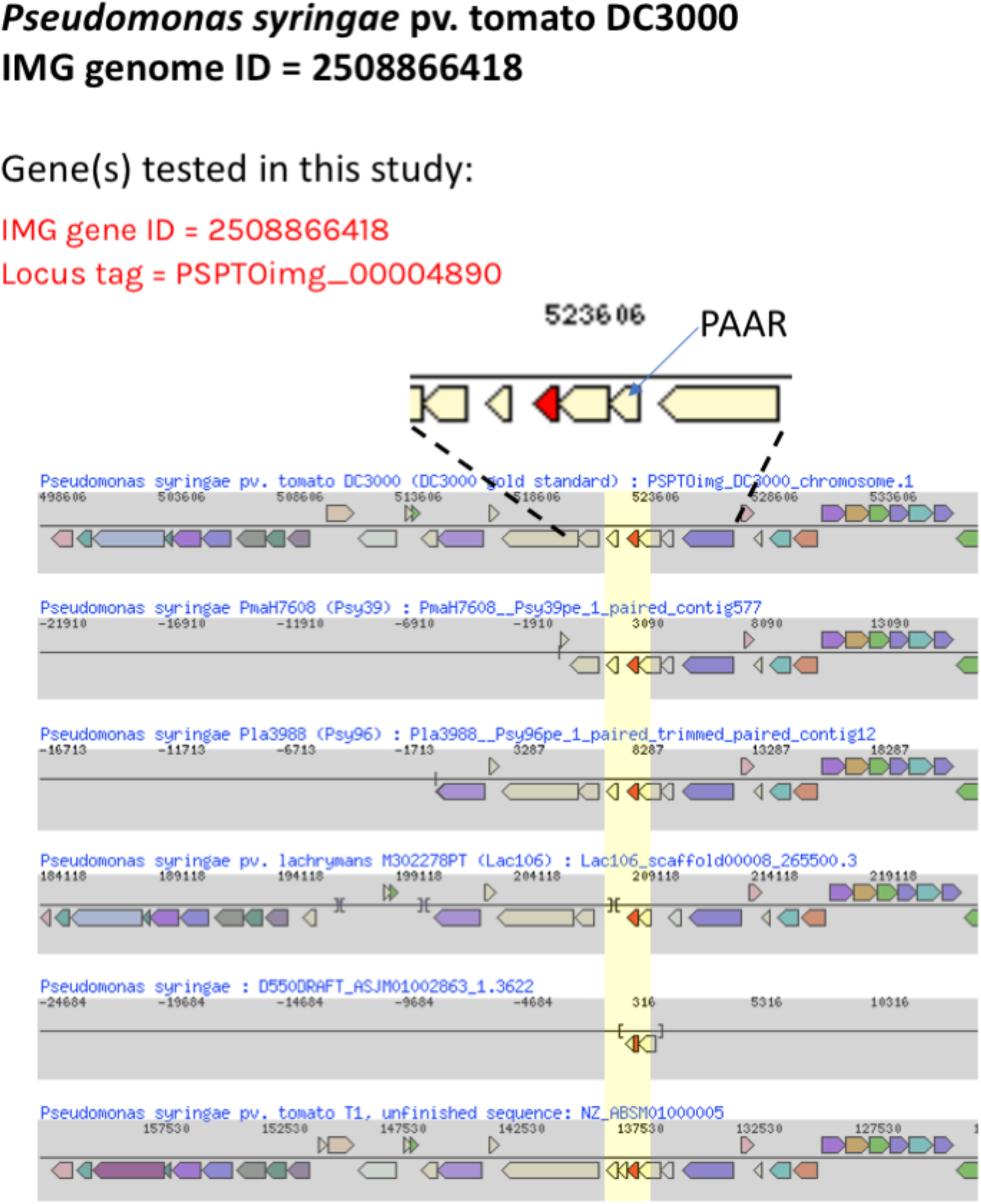
Rapid evolution of putative T6E PSPTOimg_00004890. The gene tested in this study (top, red) has a PAAR in its orphan operon. Upon looking at closely related orthologs, we see copy number variation of the red gene in the same locus (the area of duplication and deletion is highlighted in yellow).

**Supplementary Figure 6.**
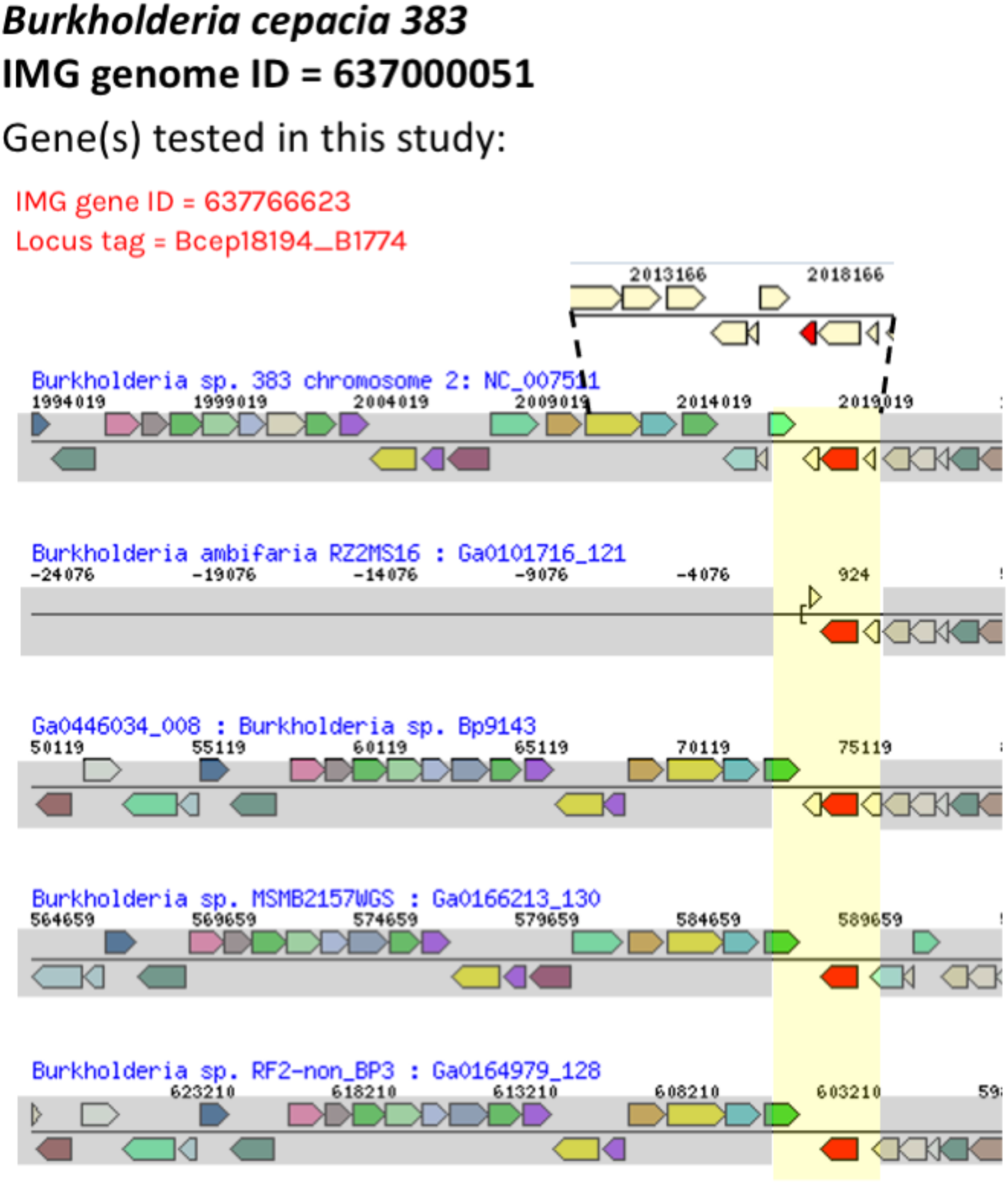
Rapid evolution of putative T6E Bcep18194_B1774. The gene tested in this study (top, red) is encoded in a “super orphan” locus, i.e. with no T6SS cores in the area. Upon looking at closely related orthologs of the adjacent putative adaptor (bottom), we see copy number variation of the putative T6E gene in the same locus (highlighted yellow).

**Supplementary Figure 7.**
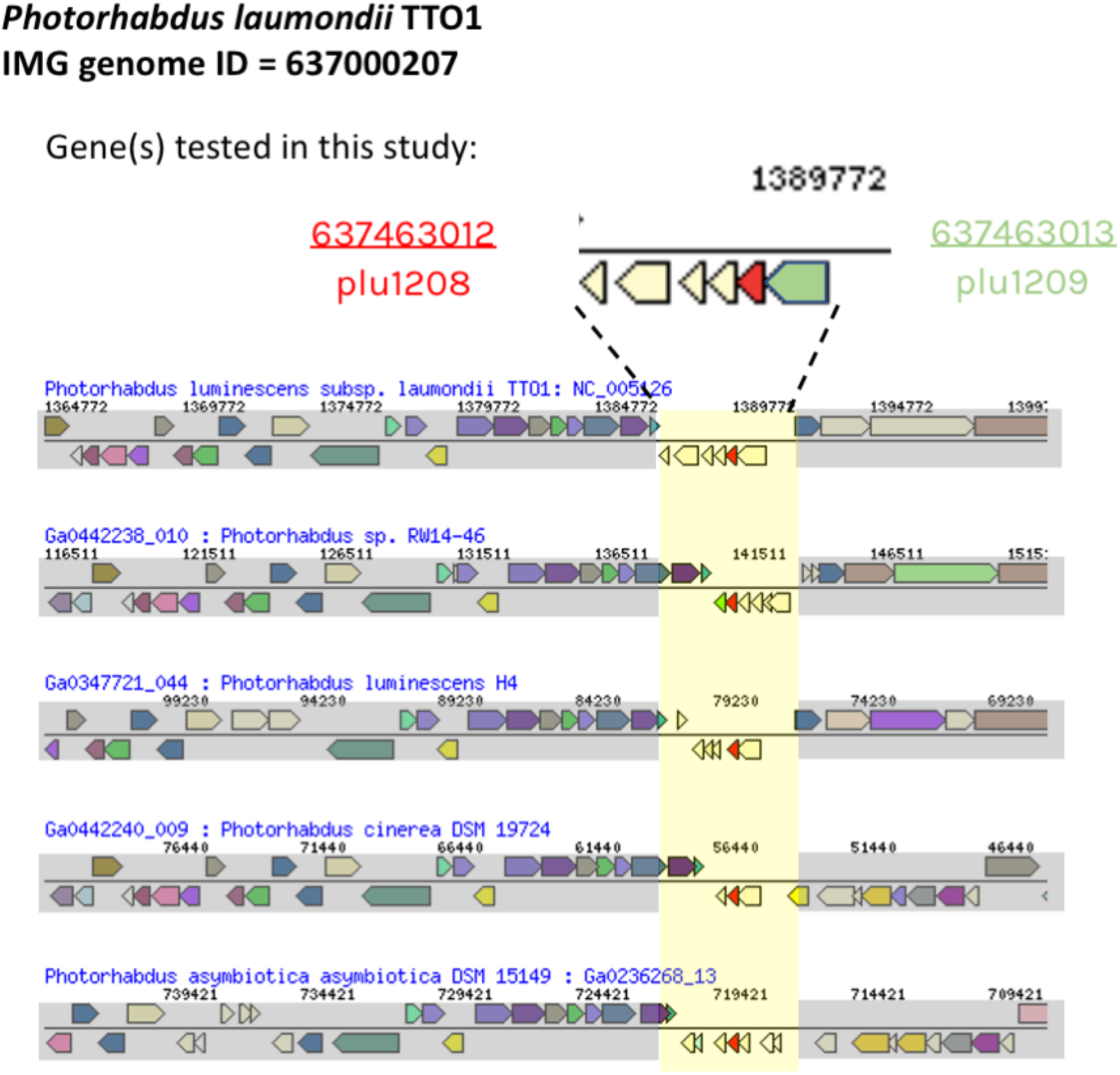
Rapid evolution of putative T6E plu1208. The genes tested in this study (top, red and green) are encoded in a “super orphan” locus, i.e. with no T6SS cores in the area. Upon looking at closely related orthologs of the adjacent putative adaptor (bottom), we see copy number variation of genes in the same locus (highlighted yellow).

**Supplementary Figure 8.**
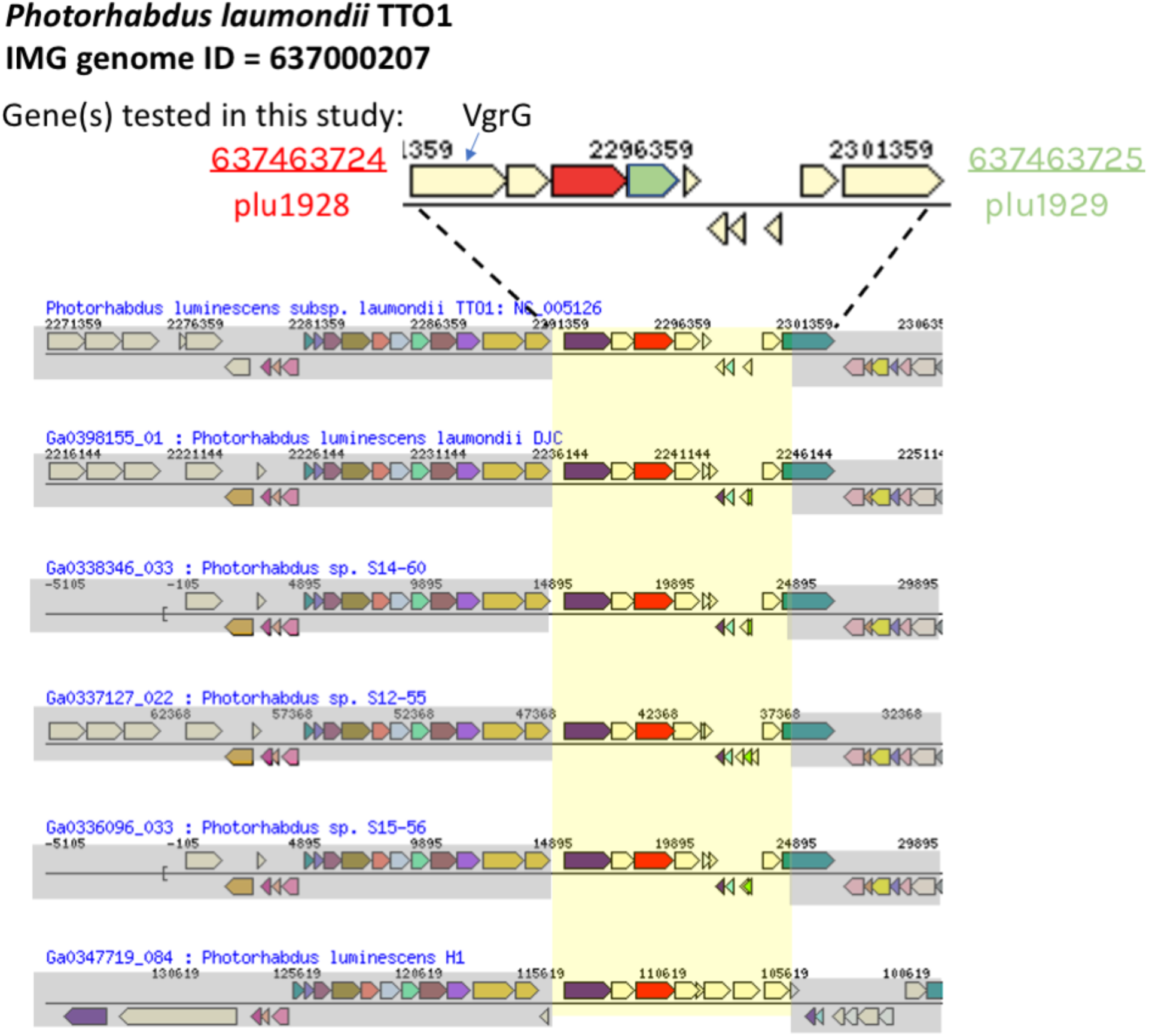
Rapid evolution of putative T6E plu1929. The genes tested in this study (top, red and green) are encoded next to a VgrG gene (T6SS core gene). Upon looking at closely related orthologs of the adjacent putative adaptor (bottom), we see copy number variation of genes in the same locus (highlighted yellow). Namely, orthologs of plu1929 can be found in up to four copies in the last genome in the row.

**Supplementary Figure 9.**
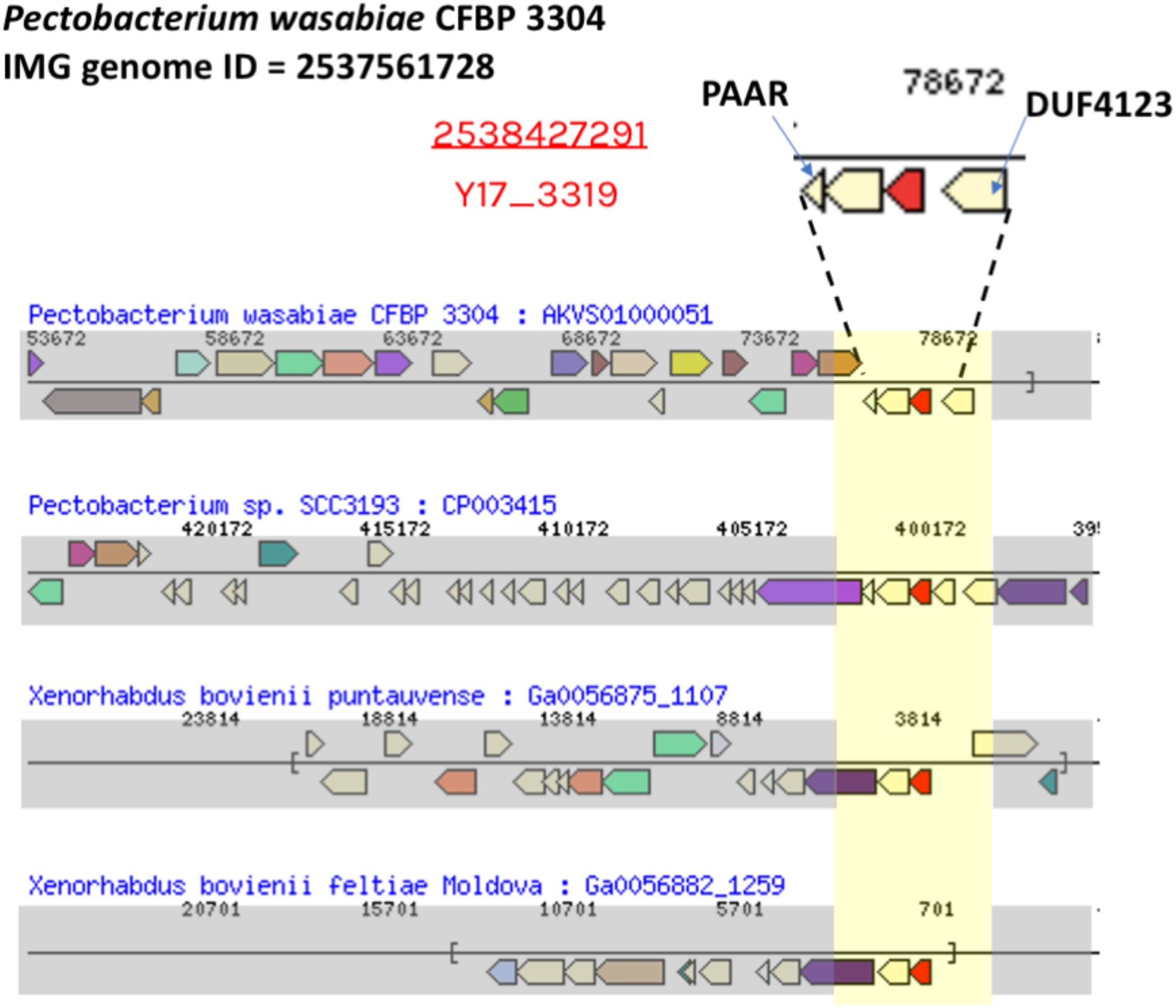
Rapid evolution of putative T6E Y17_3319. The gene tested in this study (top, red) is encoded in an orphan operon with T6SS core PAAR and known adaptor DUF4123 in the operon. Upon looking at closely related orthologs of the adjacent putative adaptor (bottom), we see copy number variation of the gene in the same locus (highlighted yellow).

**Supplementary Figure 10.**
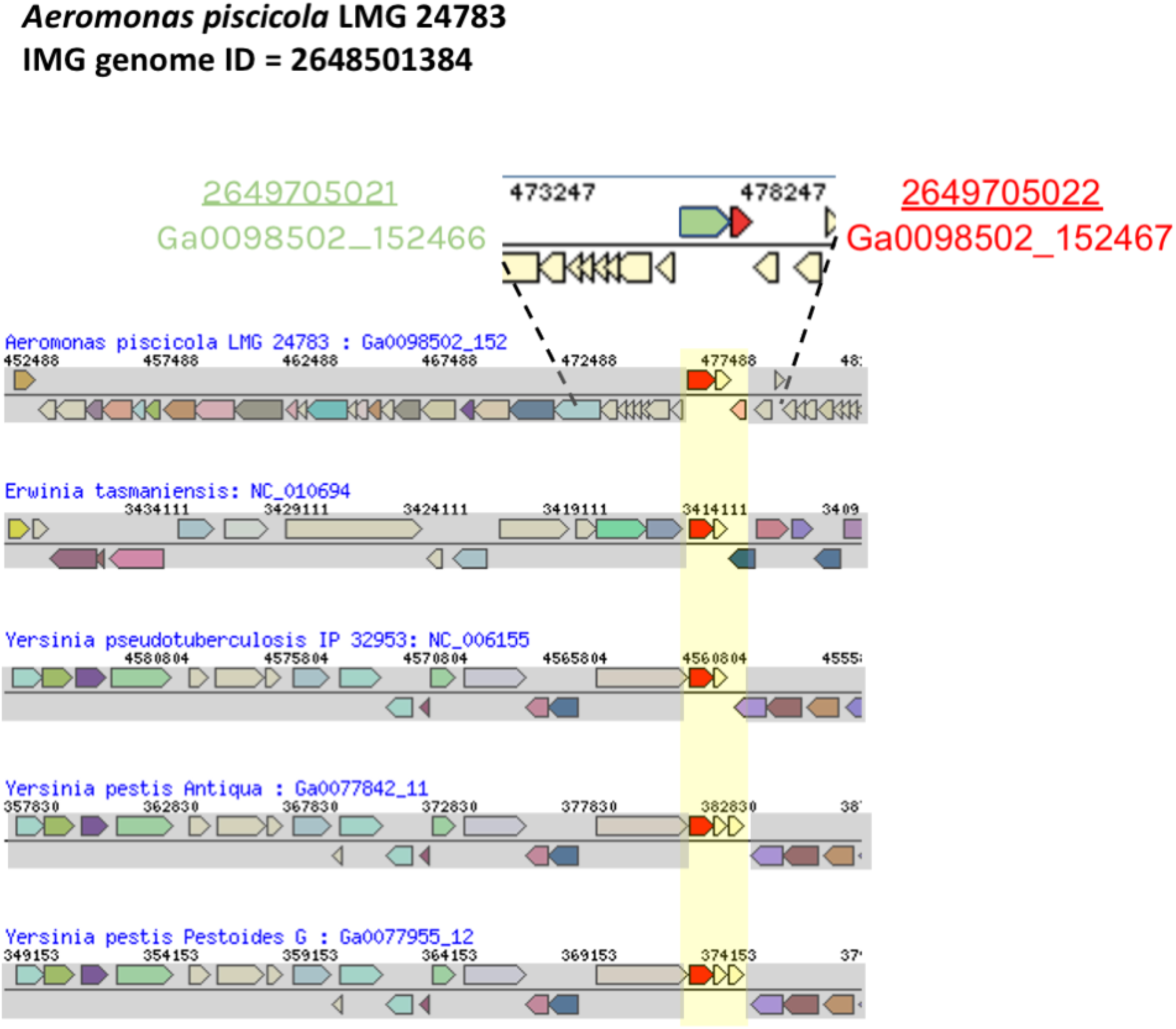
Rapid evolution of putative T6E Ga0098502_152467. The genes tested in this study (top, red and green) are encoded in a “super orphan” locus, i.e. with no T6SS cores in the area. Upon looking at closely related orthologs of the adjacent putative adaptor (bottom), we see copy number variation of genes in the same locus (highlighted yellow).

**Supplementary Figure 11.**
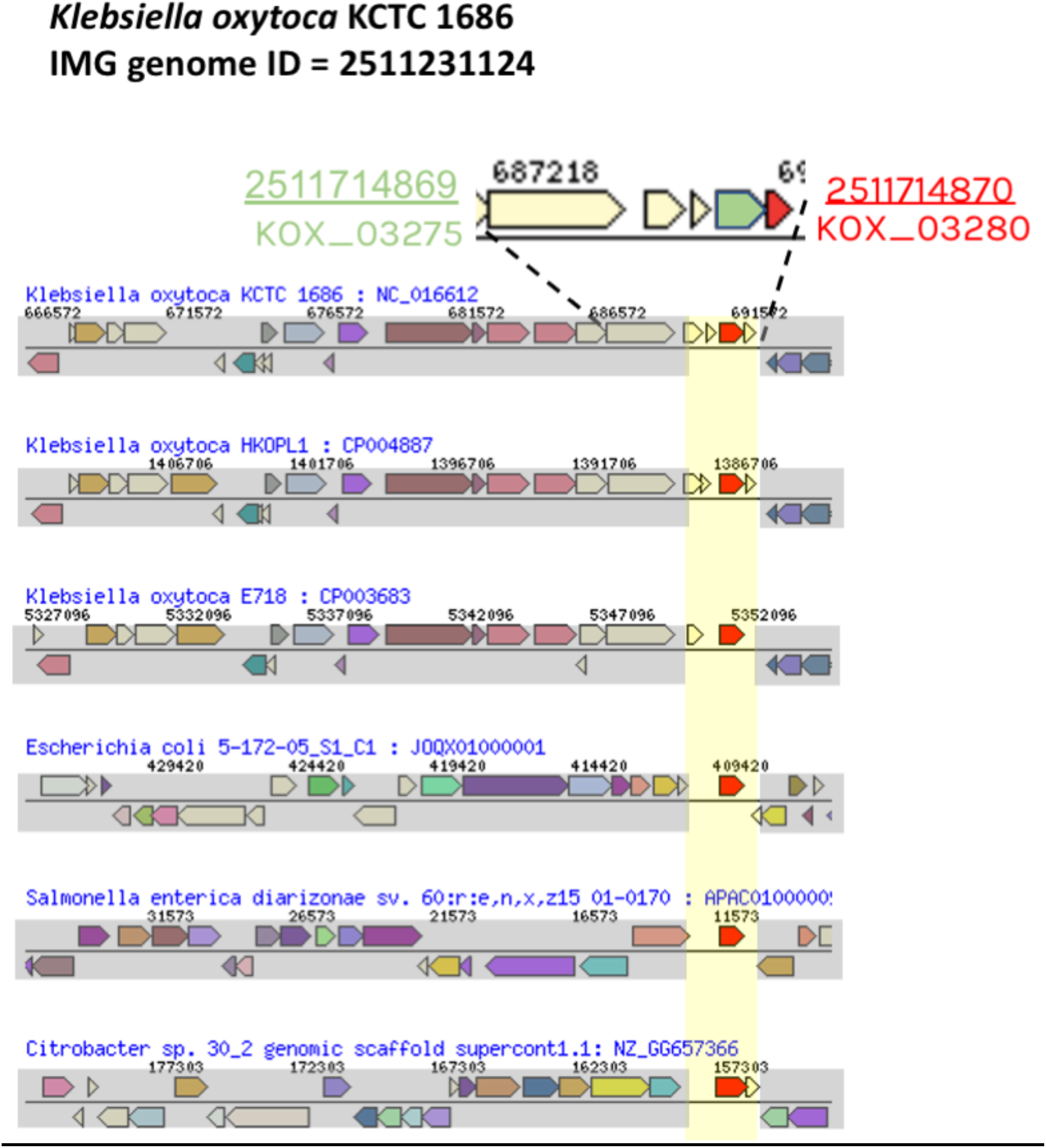
Rapid evolution of putative T6E KOX_03280. The genes tested in this study (top, red and green) are encoded in a “super orphan” locus, i.e. with no T6SS cores in the area. Upon looking at closely related orthologs of the adjacent putative adaptor (bottom), we see copy number variation of genes in the same locus (highlighted yellow). Note that both the putative adaptor and putative T6E were toxic to *E. coli*, in this case.

**Supplementary Figure 12.**
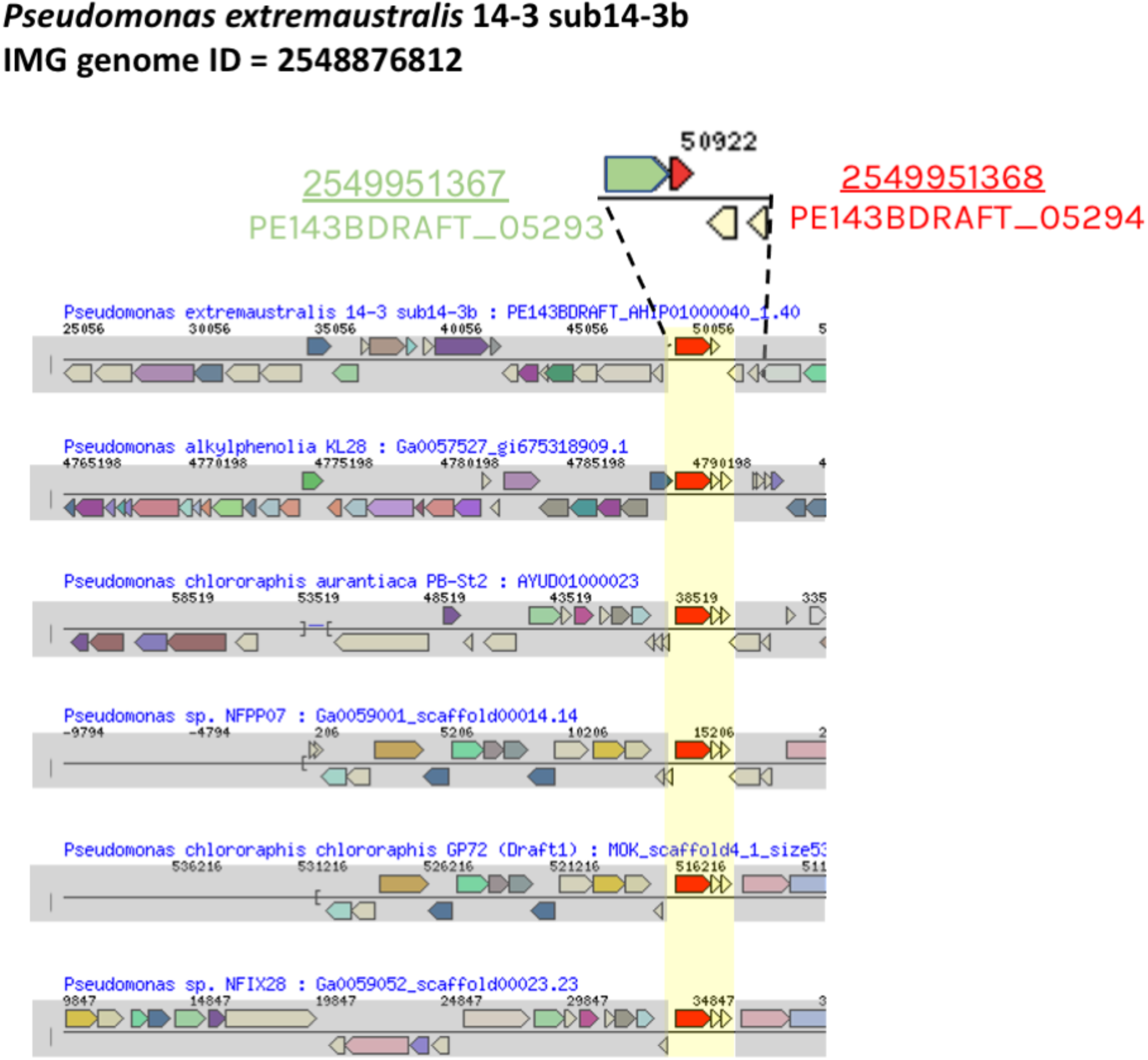
Rapid evolution of putative T6E PE143BDRAFT_05294. The genes tested in this study (top, red and green) are encoded in a “super orphan” locus, i.e. with no T6SS cores in the area. Upon looking at closely related orthologs of the adjacent putative adaptor (bottom), we see copy number variation of genes in the same locus (highlighted yellow).

**Supplementary Figure 13.**
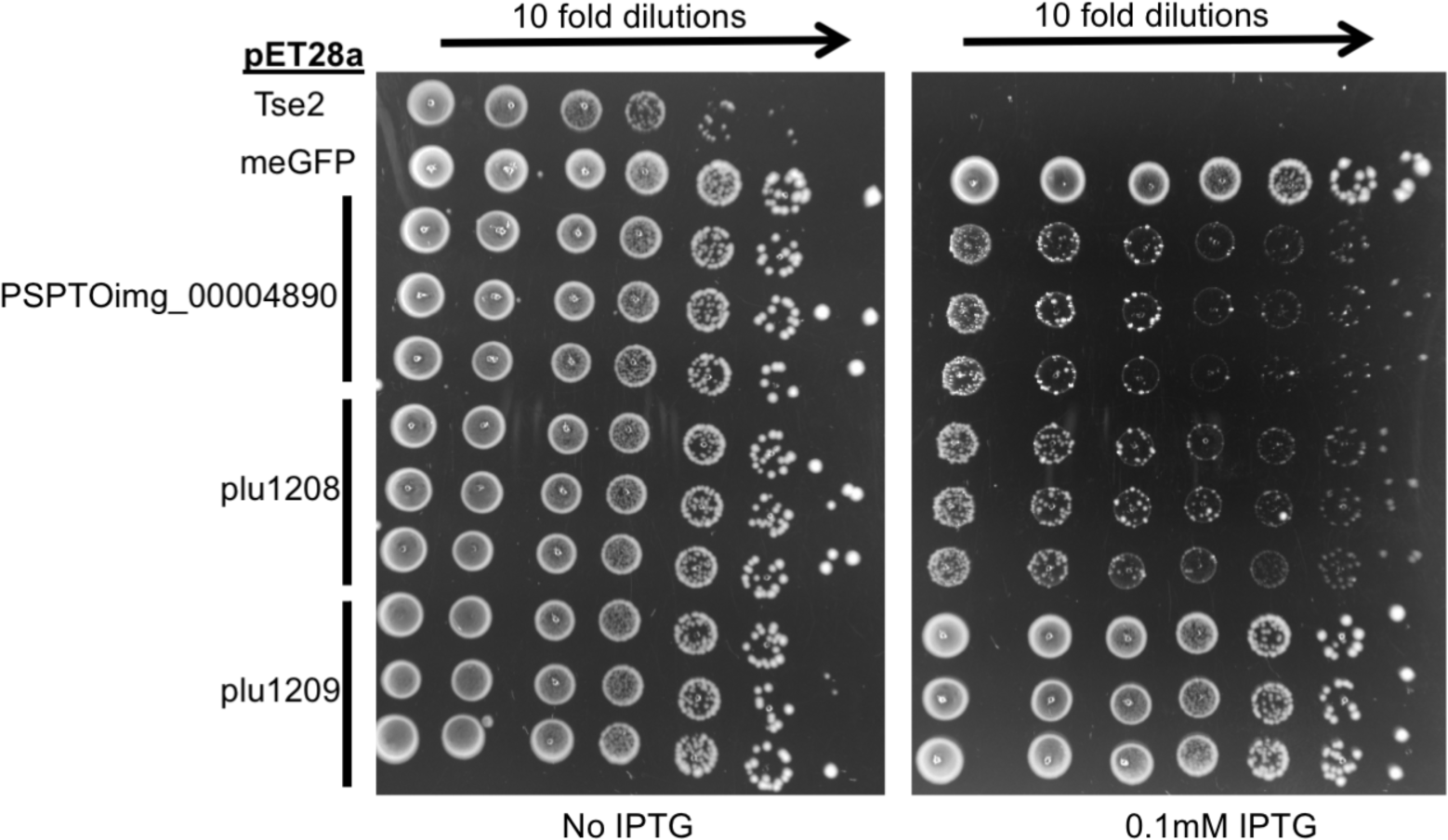
Putative T6Es PSPTOimg_00004890 and plu1208 are toxic to *E. coli*, while putative adaptor plu1209 is not. Negative controls gene mEGFP, or positive control gene, known toxin Tse2 (Hood et al. 2010), or putative T6Es and adaptors were cloned into pET28a plasmids, and transformed into *E. coli* BL21 (DE3) pLysS. Colonies were grown overnight, and were normalized and serially ten fold diluted and dropped onto agar plates in normal (left) and inducing 0.1mM IPTG (right).

**Supplementary Figure 14.**
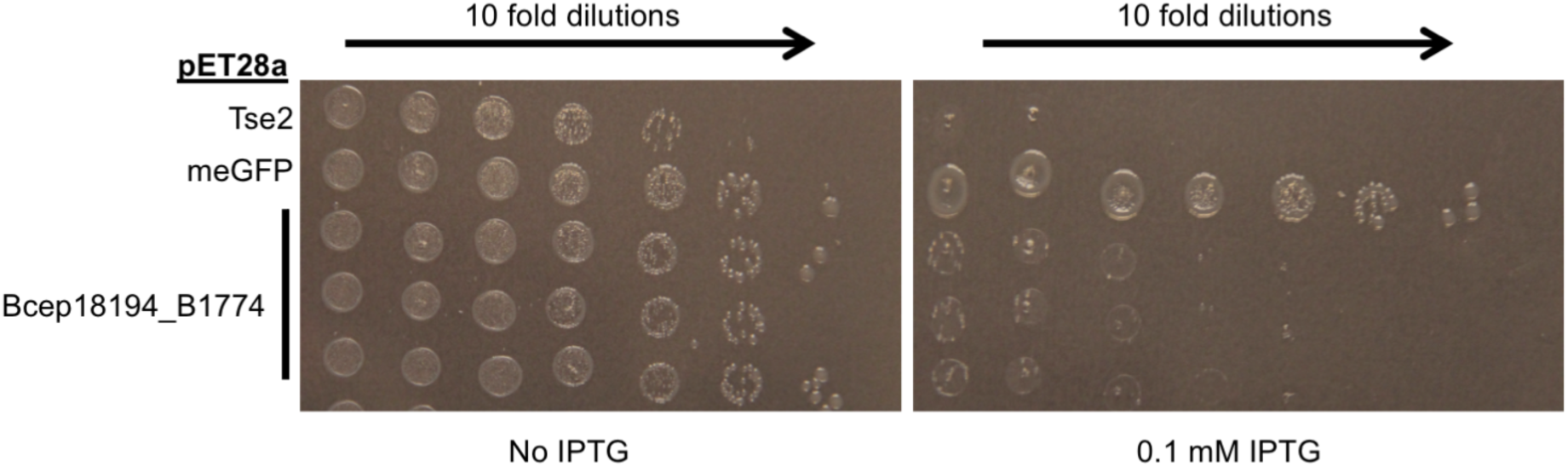
Putative T6E Bcep18194_B1774 is toxic to *E. coli*. Negative controls gene mEGFP, or positive control gene, known toxin Tse2 (Hood et al. 2010), or putative T6E Bcep18194_B1774 was cloned into pET28a plasmids, and transformed into *E. coli* BL21 (DE3) pLysS. Colonies were grown overnight, and were normalized and serially ten fold diluted and dropped onto agar plates in normal (left) and inducing 0.1mM IPTG (right).

**Supplementary Figure 15.**
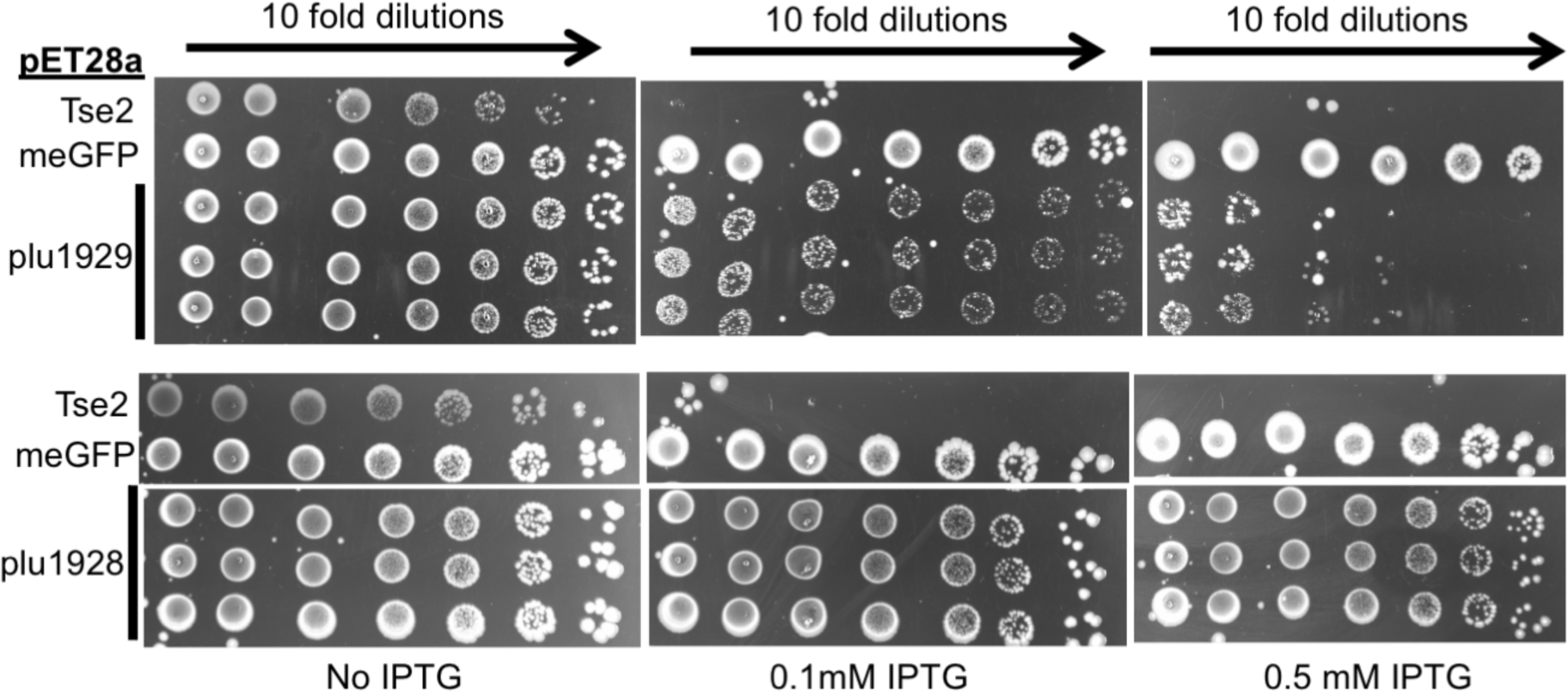
Putative T6E plu1929 is toxic to *E. coli*, while putative adaptor plu1928 is not. Negative controls gene mEGFP, or positive control gene, known toxin Tse2 (Hood et al. 2010), or putative T6E plu1929 and putative adaptor plu1928 were cloned into pET28a plasmids, and transformed into *E. coli* BL21 (DE3) pLysS. Colonies were grown overnight, and were normalized and serially ten fold diluted and dropped onto agar plates in normal (left) and inducing 0.1mM IPTG (middle), and 0.5 mM IPTG (right). This higher concentration of IPTG shows putative T6E toxicity, while maintaining putative adaptor non-toxicity, demonstrating specificity.

**Supplementary Figure 16.**
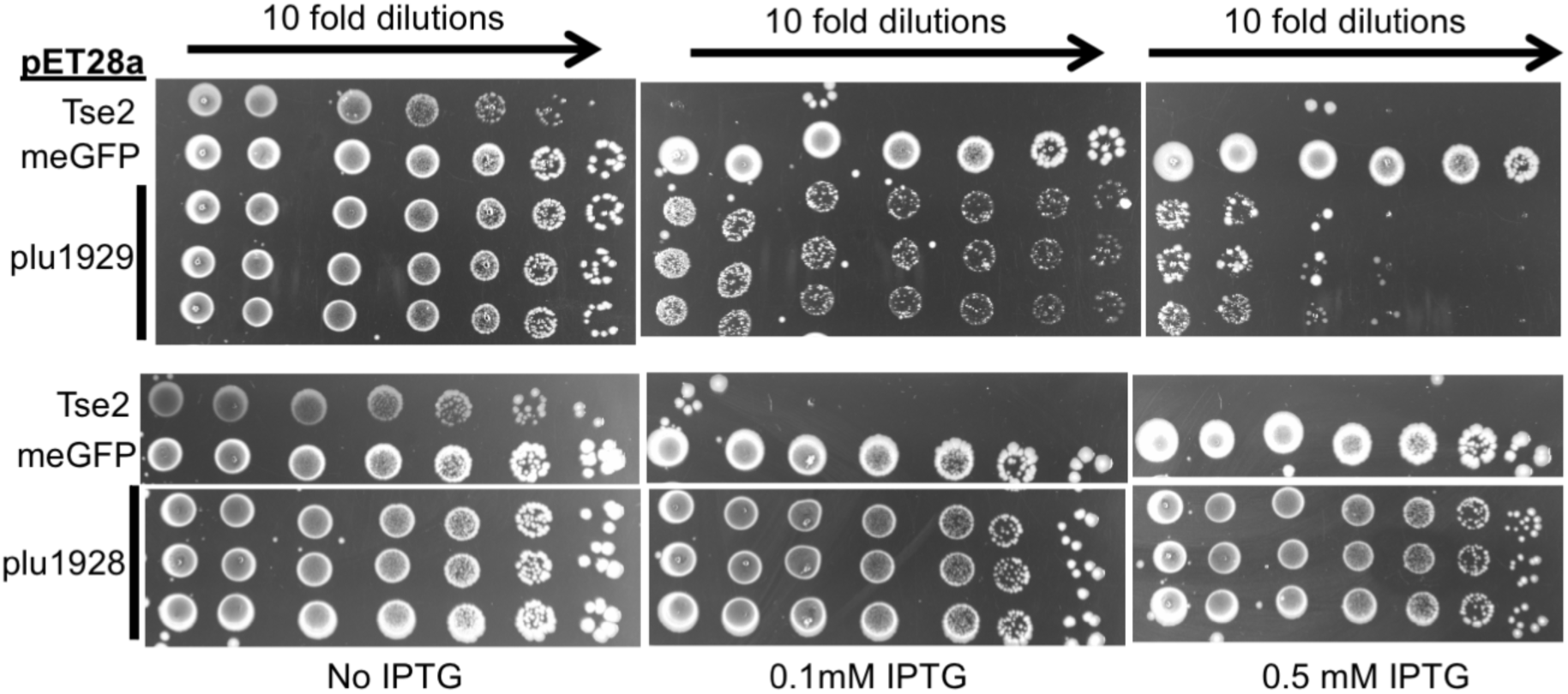
Putative T6E Y17_3319 is toxic to *E. coli*. Negative controls gene mEGFP, or positive control gene, known toxin Tse2 (Hood et al. 2010), or putative T6E Y17_3319 was cloned into pET28a plasmids, and transformed into *E. coli* BL21 (DE3) pLysS. Colonies were grown overnight, and were normalized and serially ten fold diluted and dropped onto agar plates in normal (left) and inducing 0.1mM IPTG (right).

**Supplementary Figure 17.**
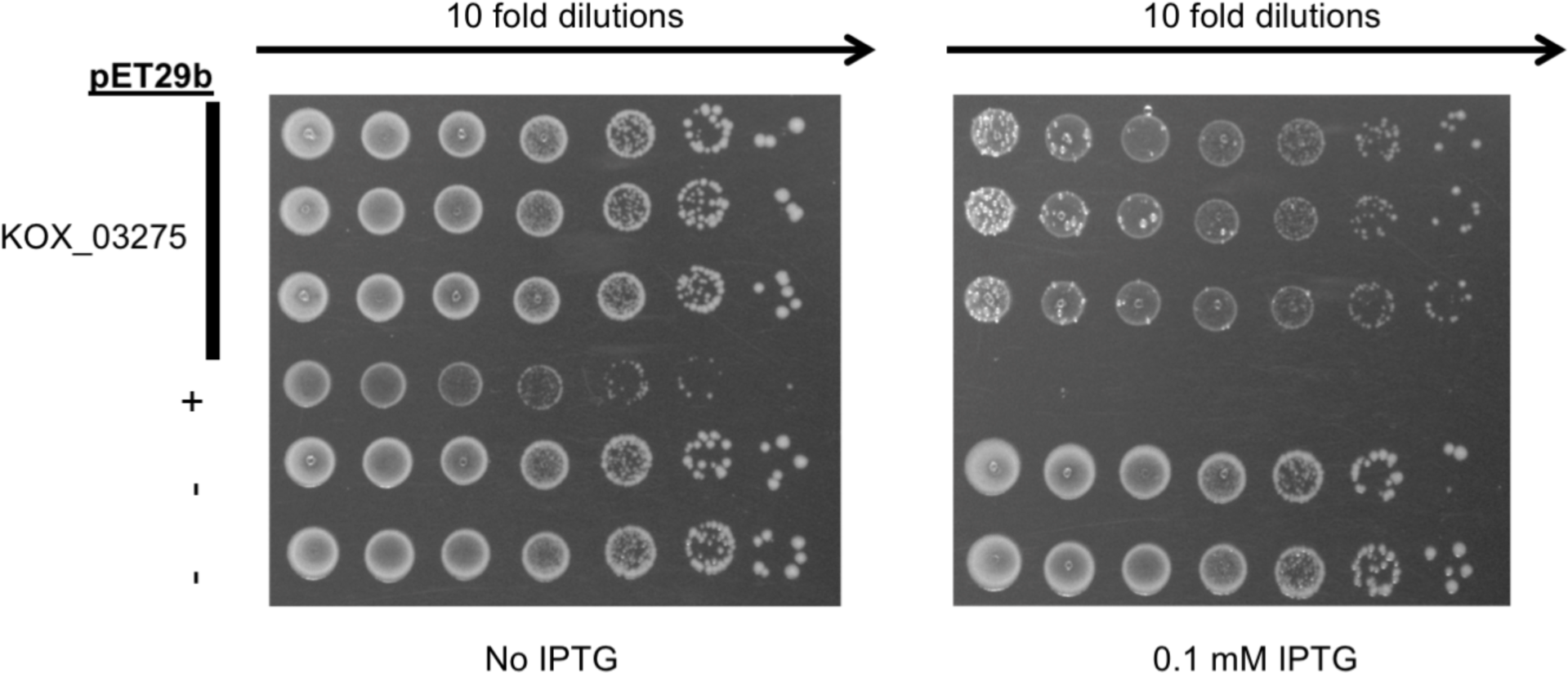
Putative adaptor KOX_03275 is toxic to *E. coli*. Empty pET29b plasmids (-), or pET29b cloned with a positive control (a known toxin, +), or putative adaptor KOX_03275 were transformed into *E. coli* BL21 (DE3). Colonies were grown overnight, and were normalized and serially ten fold diluted and dropped onto agar plates in normal (left) and inducing 0.1mM IPTG (right).

**Supplementary Figure 18.**
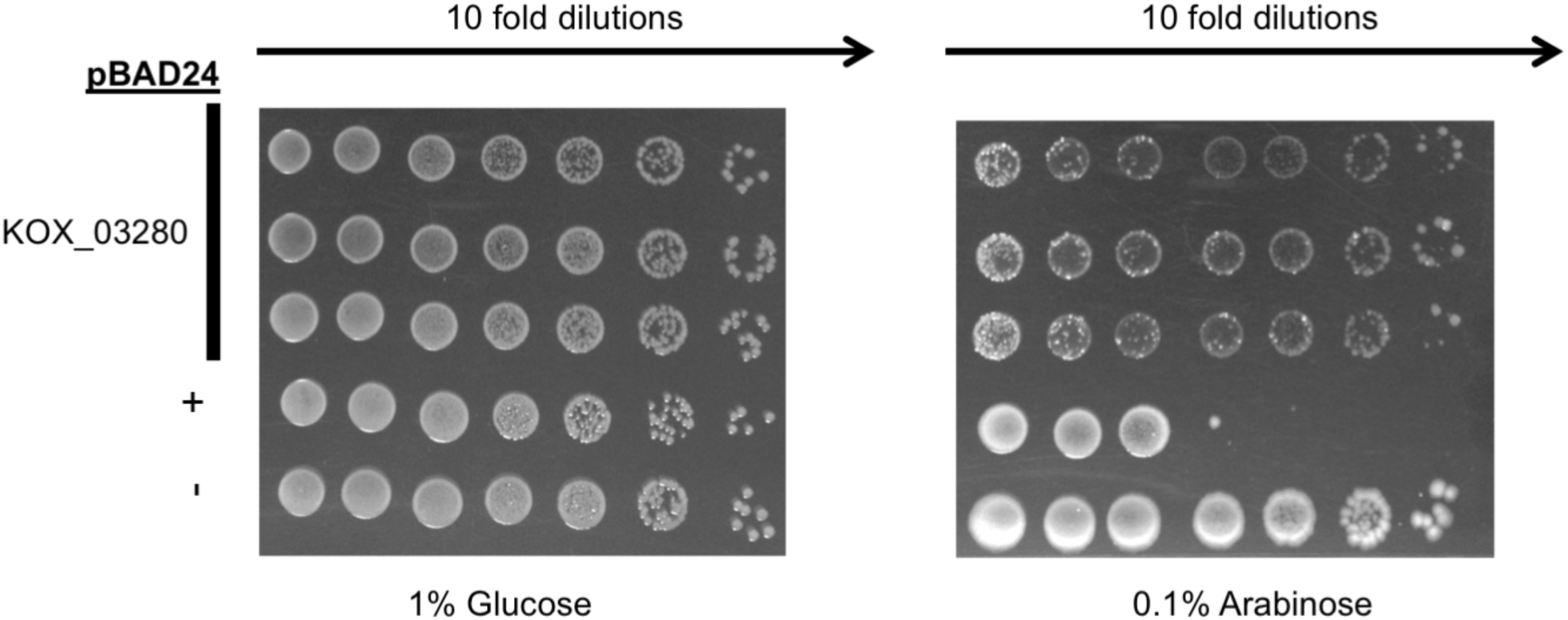
Putative T6E KOX_03280 is toxic to *E. coli*. Empty pBAD24 plasmids (-), or positive control (a known toxin, +), or putative T6E KOX_03280 was transformed into *E. coli* BL21 (DE3). Colonies were grown overnight, and were normalized and serially ten fold diluted and dropped onto agar plates with repressing 1% Glucose (left) and inducing 0.1% Arabinose (right).

**Supplementary Figure 19.**
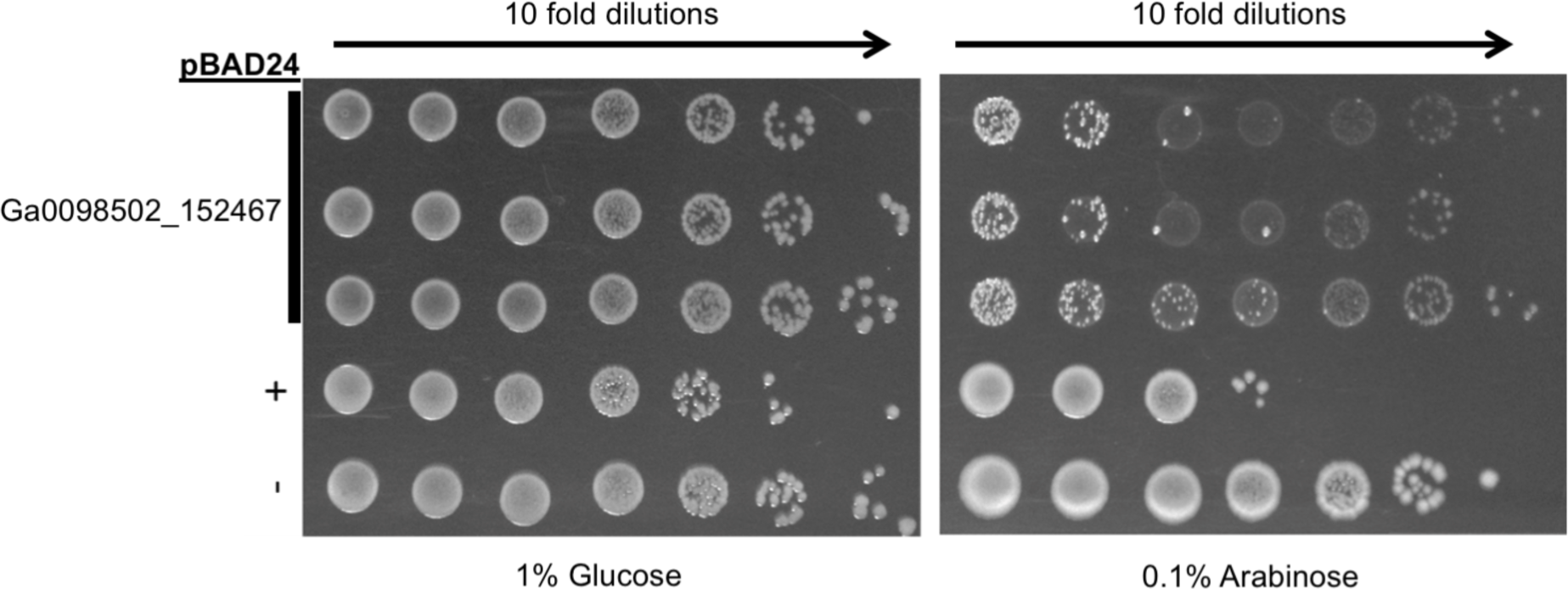
Putative T6E Ga0098502_152467 is toxic to *E. coli*. Empty pBAD24 plasmids (-), or positive control (a known toxin, +), or putative T6E Ga0098502_152467 were transformed into *E. coli* BL21 (DE3). Colonies were grown overnight, and were normalized and serially ten fold diluted and dropped onto agar plates with repressing 1% Glucose (left) and inducing 0.1% Arabinose (right).

**Supplementary Figure 20.**
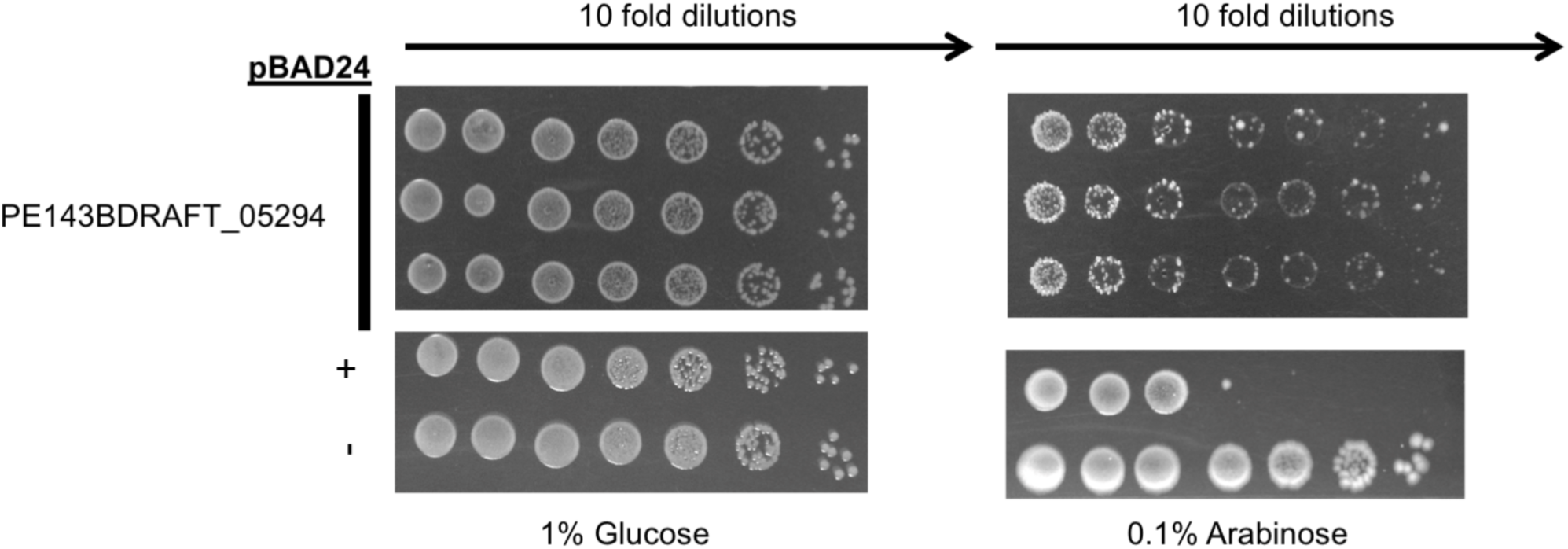
Putative T6E PE143BDRAFT_05294 is toxic to *E. coli*. Empty pBAD24 plasmids (-), or positive control (a known toxin, +), or putative T6E PE143BDRAFT_05294 were transformed into *E. coli* BL21 (DE3) pLysS. Colonies were grown overnight, and were normalized and serially ten fold diluted and dropped onto agar plates with repressing 1% Glucose (left) and inducing 0.1% Arabinose (right).

**Supplementary Figure 21.**
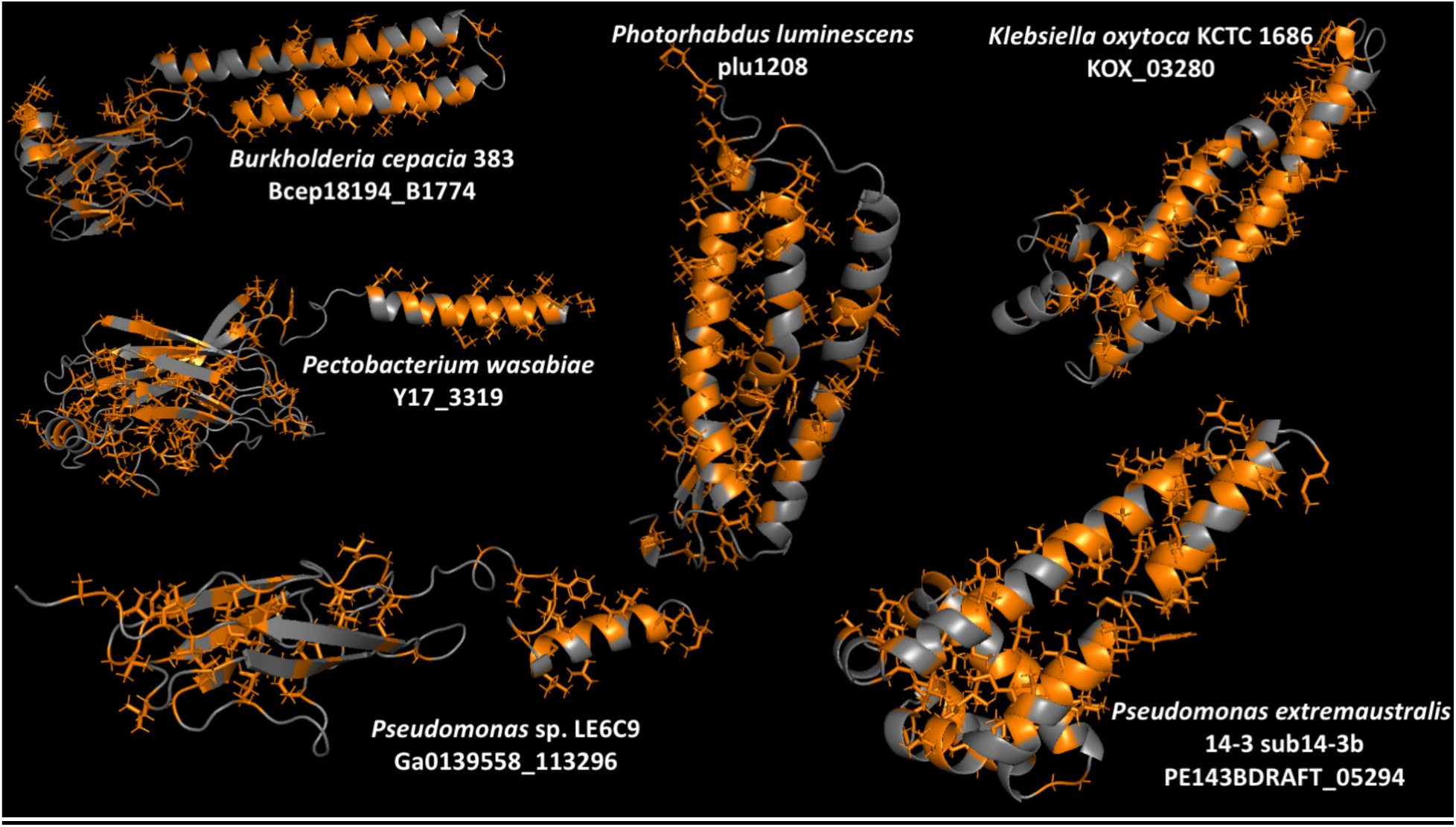
T6Es with toxic activity in *E. coli* that have hydrophobic domains. Structures predicted by Robetta’s RoseTTAFold (Baek et al. 2021) of toxic putative T6Es with transmembrane domains as determined by IMG annotation (Markowitz et al. 2012; I.-M. A. Chen et al. 2017, 2019). Hydrophobic amino acids are highlighted in orange.

**Supplementary Table 1.** Conserved T6SS domains used in this study.

| Database | Accession | Domain name | Used for |
| --- | --- | --- | --- |
| pfam | pfam05488 | PAAR domain | search for “evolved” cores |
| pfam | pfam13665 | DUF4150 (PAAR like) | search for “evolved” cores |
| pfam | pfam05954 | Phage GPD protein (VgrG analog) | search for “evolved” cores |
| pfam | pfam06715 | gp5 repeats (VgrG component) | search for “evolved” cores |
| pfam | pfam04717 | base V (VgrG component) | search for “evolved” cores |
| pfam | pfam13296 | T6SS_VgrG | search for “evolved” cores |
| pfam | pfam10106 | DUF2345 | search for “evolved” cores |
| pfam | pfam05638 | HCP | search for “evolved” cores |
| pfam | pfam17642 | TssD | search for “evolved” cores |
| COG | COG3455 | VasL (TssL) | Defining T6SS+ genomes |
| COG | COG3521 | TssJ | Defining T6SS+ genomes |
| COG | COG3523 | TssM | Defining T6SS+ genomes |

**Supplementary Table 2.**
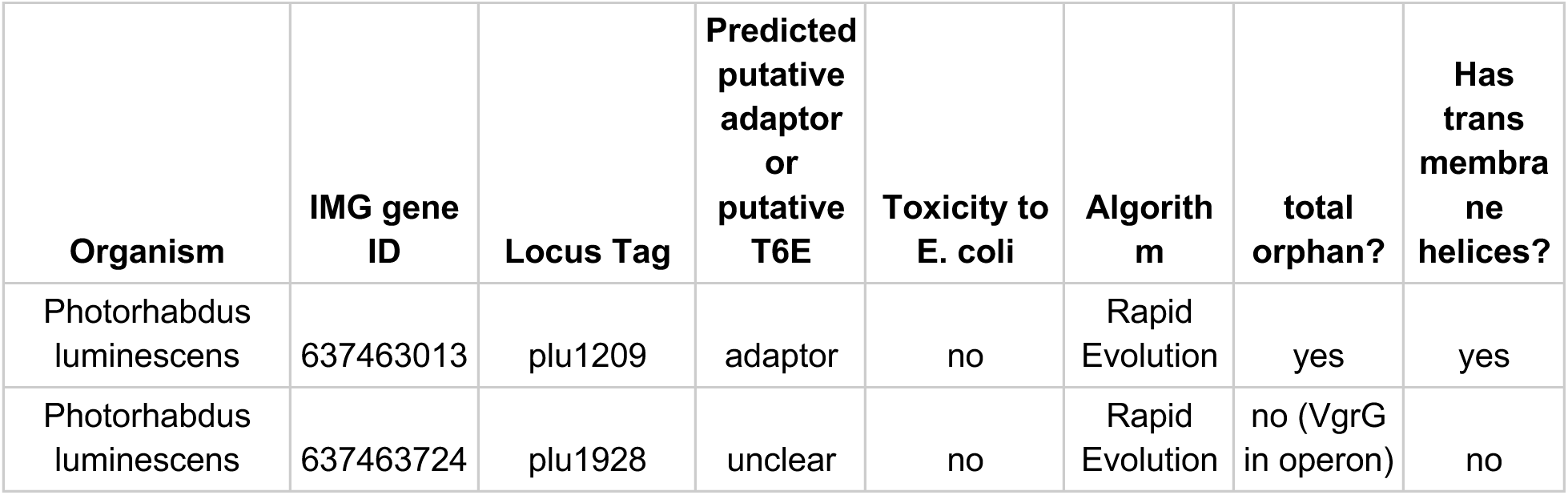
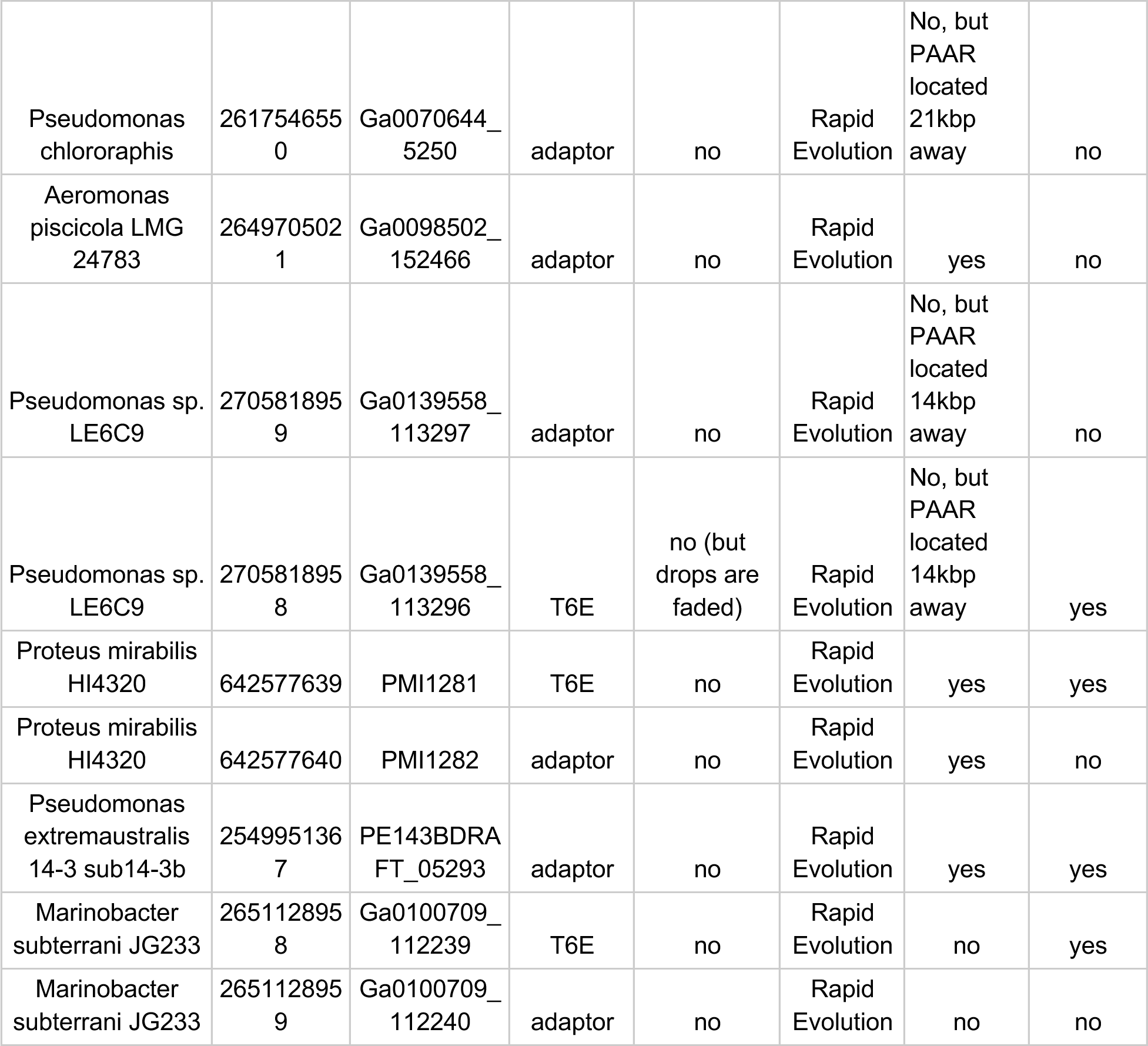
Tested genes that were not toxic to *E. coli*.

